# Network analysis reveals strain-dependent response to misfolded tau aggregates

**DOI:** 10.1101/2023.01.28.526029

**Authors:** Dominic J. Acri, Yanwen You, Mason D. Tate, Brianne McCord, A. Daniel Sharify, Sutha John, Hande Karahan, Byungwook Kim, Luke C. Dabin, Stéphanie Philtjens, H.R. Sagara Wijeratne, Tyler J. McCray, Daniel C. Smith, Stephanie J. Bissel, Bruce T. Lamb, Cristian A. Lasagna-Reeves, Jungsu Kim

**Affiliations:** Stark Neurosciences Research Institute, Indiana University School of Medicine, Indianapolis, IN 46202, USA; Medical Neuroscience Graduate Program, Indiana University School of Medicine, Indianapolis, IN 46202, USA; Department of Anatomy, Cell Biology & Physiology, Indiana University School of Medicine, Indianapolis, IN, USA; Department of Medical and Molecular Genetics, Indiana University School of Medicine, Indianapolis, IN 46202, USA; Department of Biochemistry and Molecular Biology, Indiana University School of Medicine, Indianapolis, IN 46202, USA; Center for Computational Biology and Bioinformatics, Indiana University School of Medicine, Indianapolis, IN, USA

## Abstract

Mouse genetic backgrounds have been shown to modulate amyloid accumulation and propagation of tau aggregates. Previous research into these effects has highlighted the importance of studying the impact of genetic heterogeneity on modeling Alzheimer’s disease. However, it is unknown what mechanisms underly these effects of genetic background on modeling Alzheimer’s disease, specifically tau aggregate-driven pathogenicity. In this study, we induced tau aggregation in wild-derived mice by expressing *MAPT* (P301L). To investigate the effect of genetic background on the action of tau aggregates, we performed RNA sequencing with brains of 6-month-old C57BL/6J, CAST/EiJ, PWK/PhJ, and WSB/EiJ mice (n=64). We also measured tau seeding activity in the cortex of these mice. We identified three gene signatures: core transcriptional signature, unique signature for each wild-derived genetic background, and tau seeding-associated signature. Our data suggest that microglial response to tau seeds is elevated in CAST/EiJ and PWK/PhJ mice. Together, our study provides the first evidence that mouse genetic context influences the seeding of tau.

**Graphical Abstract:** 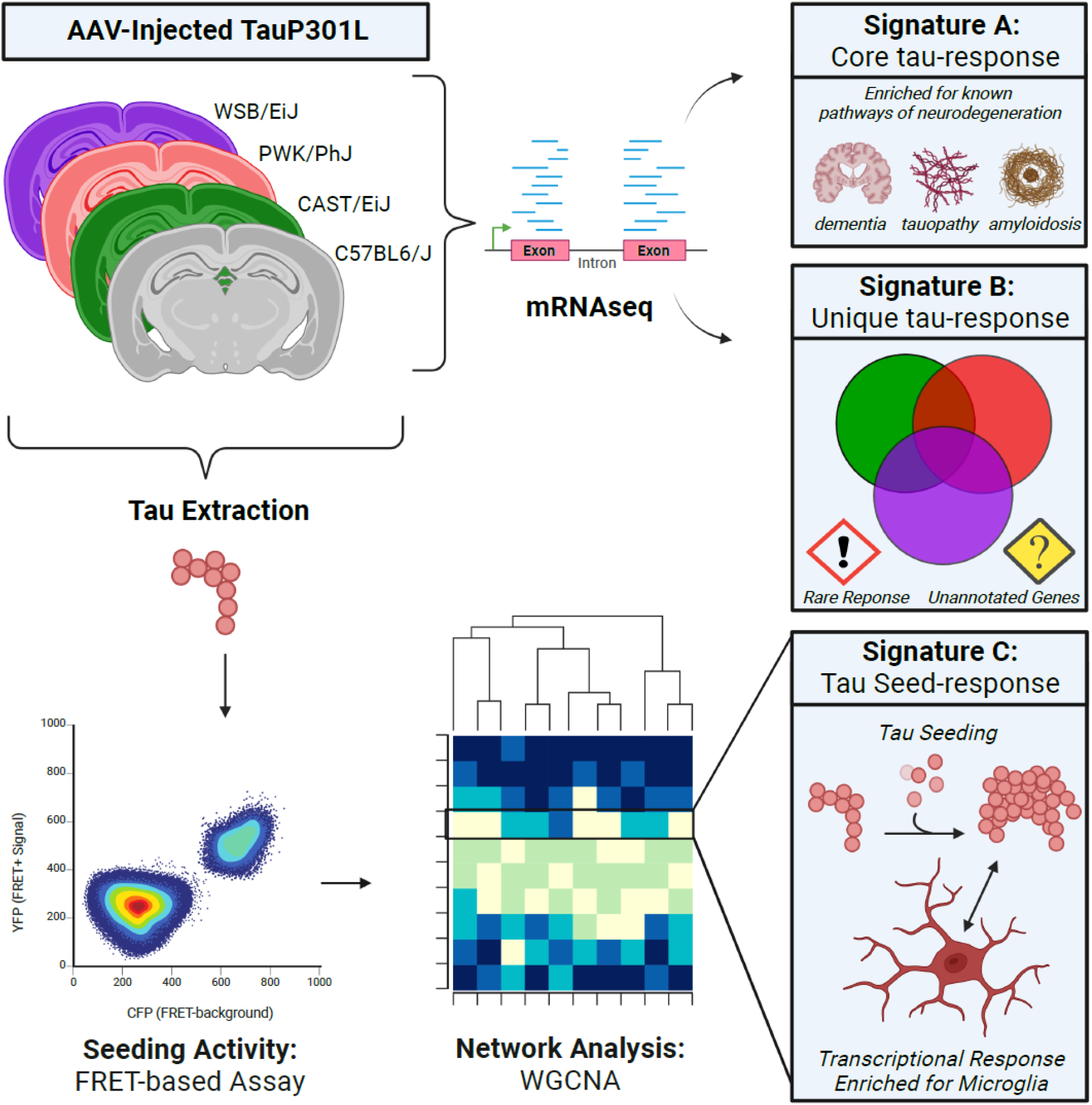

**SUMMARY:** Seeding of tau predates the phosphorylation and spreading of tau aggregates. Acri and colleagues report transcriptomic responses to tau and elevated tau seeds in wild-derived mice. This paper creates a rich resource by combining genetics, tau biosensor assays, and transcriptomics.

## INTRODUCTION

Alzheimer’s disease (AD) is the most common cause of dementia and is characterized by the accumulation of amyloid plaques and neurofibrillary tangles mainly comprised of aggregated tau protein (Long and Holtzman, 2019). Human genetic studies have identified variants that implicate several risk genes that influence AD pathogenesis (Karch and Goate, 2015). As researchers design studies to investigate the role of these late-onset AD risk genes, they must first decide which pathological outcome(s) to measure. A combination of transgenic, viral, and xenograft approaches have been developed to study amyloid-only, tau-only, and amyloid-tau pathogenesis in mice. While the ultimate goal of these studies is to translate findings to patients with AD, the first step to translation is understanding what is fundamentally happening in the model organism.

There are a number of promising therapeutic approaches that target tau (Congdon and Sigurdsson, 2018). Importantly, hyperphosphorylated tau has been shown to cause neuronal cell death (Lee et al., 2011) and to correlate with measures of cognitive decline (Arriagada et al., 1992) in humans. To investigate the progression of tauopathy, the most widely used mouse models express Microtubule Associated Protein Tau (*MAPT*), the gene which encodes the tau protein. The P301L mutation originally described in frontotemporal dementia patients (Hutton et al., 1998, Poorkaj et al., 1998) is often used to induce tau aggregate formation and study tau pathogenesis. Transgenic models of Tau^P301L/^S(Santacruz et al., 2005, Yoshiyama et al., 2007) and viral models of Tau^P301L^ (Cook et al., 2015, Wegmann et al., 2017) are useful tools to study the progression of tau pathology and investigate factors that could lead to the risk of developing any tauopathy, including AD. These models have been shown to recapitulate key aspects of human tauopathy including behavioral deficits (Lasagna-Reeves et al., 2016, Cook et al., 2014), neuroinflammation (Yoshiyama et al., 2007), prion-like proteopathic seeding (Martinez et al., 2022), and propagation from one cell to another (Dujardin et al., 2022, Rauch et al., 2020, Wegmann et al., 2017, de Calignon et al., 2012, Woerman et al., 2017).

Although the exact mechanism by which tau aggregates form is currently unknown, there is strong evidence for the role of “tau seeding” as an initiating event. Proteopathic tau seeds are capable of entering a cell and promoting aggregation in a prion-like manner (Clavaguera et al., 2009, Frost et al., 2009). Several studies have shown that seeding precedes tau pathogenesis and can even occur in brain regions where tau pathology does not usually present (DeVos et al., 2018, Kaufman et al., 2018, Stopschinski et al., 2021). Several *in vitro* models have been developed that can measure tau seeding activity from human patients or mouse models of tauopathy (Bengoa-Vergniory et al., 2021, Holmes et al., 2014, Jin et al., 2022). These seeding activity assays have assisted in the discovery of novel tau interactors and been used to investigate phosphorylation patterns associated with tau progression (Martinez et al., 2022, Mirbaha et al., 2022). Unlike human patients who are genetically diverse, most studies use the same monogenic mouse models. Therefore, the influence of genetic diversity on tau pathology and seeding has not been thoroughly investigated. With the hope that these preclinical studies will translate to tau-targeted treatments, there is a need to better understand how the genetic context of our mouse models affects our interpretation of tauopathy.

The most widely used mouse strain in biomedical research, the C57BL/6J strain (herein referred to as B6), was established by the Jackson laboratory in the 1920s and became the strain used to create the mouse reference genome (Mekada et al., 2009, Mouse Genome Sequencing et al., 2002). While one goal of sustaining a single inbred line is to limit inter-laboratory artifacts, research into B6 mice reveals that genetic drift and mixed background breeding have introduced a number of variants since the first draft of the mouse reference genome (Sarsani et al., 2019, Simon et al., 2013). These variants and others purposefully introduced by selective breeding are termed “genetic diversity.” Unique phenotypes arising from mouse genetic diversity can be used as a model for complex diseases. For example, decreases in pancreatic insulin at 12 weeks of age in the NOD/ShiLtJ mouse model established this strain as the leading model for research in Type 1 Diabetes (Makino et al., 1980). Another key strategy in harnessing mouse genetic diversity is to breed together different mouse strains to create multiparent panels for genetic mapping (Churchill et al., 2004, Churchill et al., 2012, Peirce et al., 2004).

The founder strains of the Jackson laboratory’s multiparent panels, the Diversity Outbred and Collaborative Cross mice, include 5 classically inbred and 3 wild-derived mouse strains (Churchill et al., 2012). These eight founders were selected as they could be bred together to contain segregating variants every 100-200 base pairs (Churchill et al., 2004). The most genetically distinct of the eight founder strains are the wild-derived: CAST/EiJ, PWK/PhJ, and WSB/EiJ (herein referred to as CAST, PWK, and WSB). These three wild-derived strains are descendants of three different subspecies of *Mus musculus* and contain millions of variants relative to the mouse reference genome (Yang et al., 2011). For this reason, wild-derived mouse strains have been used as a resource for modeling the population-level heterogeneity that cannot be investigated using classical inbred mouse strains alone. Deep characterization of these wild-derived mice has uncovered genetic (Morgan et al., 2015), behavioral (Kollmus et al., 2020), and immune (Lilue et al., 2018) differences that are improving our knowledge of mouse genetics.

Previous research has demonstrated the importance of studying these wild-derived mice in the context of AD. Mice with APP^swe^ and PSEN1^de9^ transgenes (*APP/PS1* transgenic: B6.Cg-Tg(Appswe,PSEN1dE9)85Dbo/Mmjax) were backcrossed onto each of these three wild-derived mouse backgrounds. These mice had higher levels of Amyloid-β (Aβ) compared to age-matched mice on a B6 background (Onos et al., 2019). Notably, Onos and colleagues observed an increase in neuroinflammation in the cortex and hippocampus of PWK mice. This is especially interesting given the important role of neuroinflammation in Aβ accumulation in AD (Efthymiou and Goate, 2017, Karahan et al., 2021, Mhatre et al., 2015, Schoch et al., 2021). Single cell RNA sequencing of sorted microglia further demonstrated that the responses of immune cell subtypes, namely homeostatic microglia and disease-associated microglia (DAMs), are determined by wild-derived genetic backgrounds (Yang et al., 2021). As the increasing focus is spent on defining microglial subtypes in studies of neurodegeneration (Keren-Shaul et al., 2017, Paolicelli et al., 2022), the effect of wild-derived genetic background could be an important factor in selecting a mouse model that better reflects human disease. Even though wild-derived backgrounds have been shown to have a large effect on modeling Aβ pathology, little is known about their effect on modeling tauopathy.

Given the known effect of wild-derived mice on modeling Aβ accumulation, we aimed to investigate the effect of wild-derived mouse genetic background on tauopathy. To preserve the mouse genetic background, we expressed mutant tau with the P301L mutation in the brains of B6, CAST, PWK, and WSB mice using intracerebroventricular injection of an adeno-associated virus (AAV) (Carlomagno et al., 2019, Cook et al., 2015). This strategy allows us to change the genetic background without the need to backcross a conventional transgenic mouse with each strain for 10+ generations. In addition to saving time and money using our viral approach, most importantly, we ensure that each genetic background is preserved without any possible genetic drift or the addition of unwanted variants over a multi-year backcrossing experiment. We found that the presence of seed-competent tau was modulated by genetic background, independent of human tau expression level. Using bulk mRNA sequencing, we report transcriptional changes that are shared across genetic backgrounds, changes that are unique to wild-derived mice, and changes that are associated with the presence of seed-competent tau (Signatures A, B, and C respectively). Our data serve as a resource for those studying the pathogenesis of tau and implicate several transcriptional signatures that are not present when modeling tauopathy in B6 mice.

## RESULTS

### Variants in Wild-derived Genetic Backgrounds within AMP-AD Nominated Target Genes

CAST, PWK, and WSB mice contain millions of variants across the mouse genome (Blake et al., 2021). These variants include over 5 million single nucleotide polymorphisms (SNPs) (Onos et al., 2019), 116 novel genes not present in B6 (Lilue et al., 2018), and between 250-400 large structural variants (Yalcin et al., 2011). To identify genetic variants in our mice, we used the Illumina Infinium platform, containing 143,259 probes, designed specifically for wild-derived mice and other founders of the Diversity Outbred mouse model (Morgan et al., 2015). This allowed us to confirm the genotype of each genetic background in our laboratory and gave us information about the SNPs and Copy Number Variants (CNVs) in structurally polymorphic regions of the mouse genome. To determine which of these variants could be important in AD research, we focused on target genes nominated by the Accelerating Medicines Partnership Program for Alzheimer’s Disease (AMP-AD) consortium. The genotyped variants of CAST, PWK, and WSB within the AMP-AD nominated targets were visualized using a circos plot (Figure 1).

**Figure 1.**
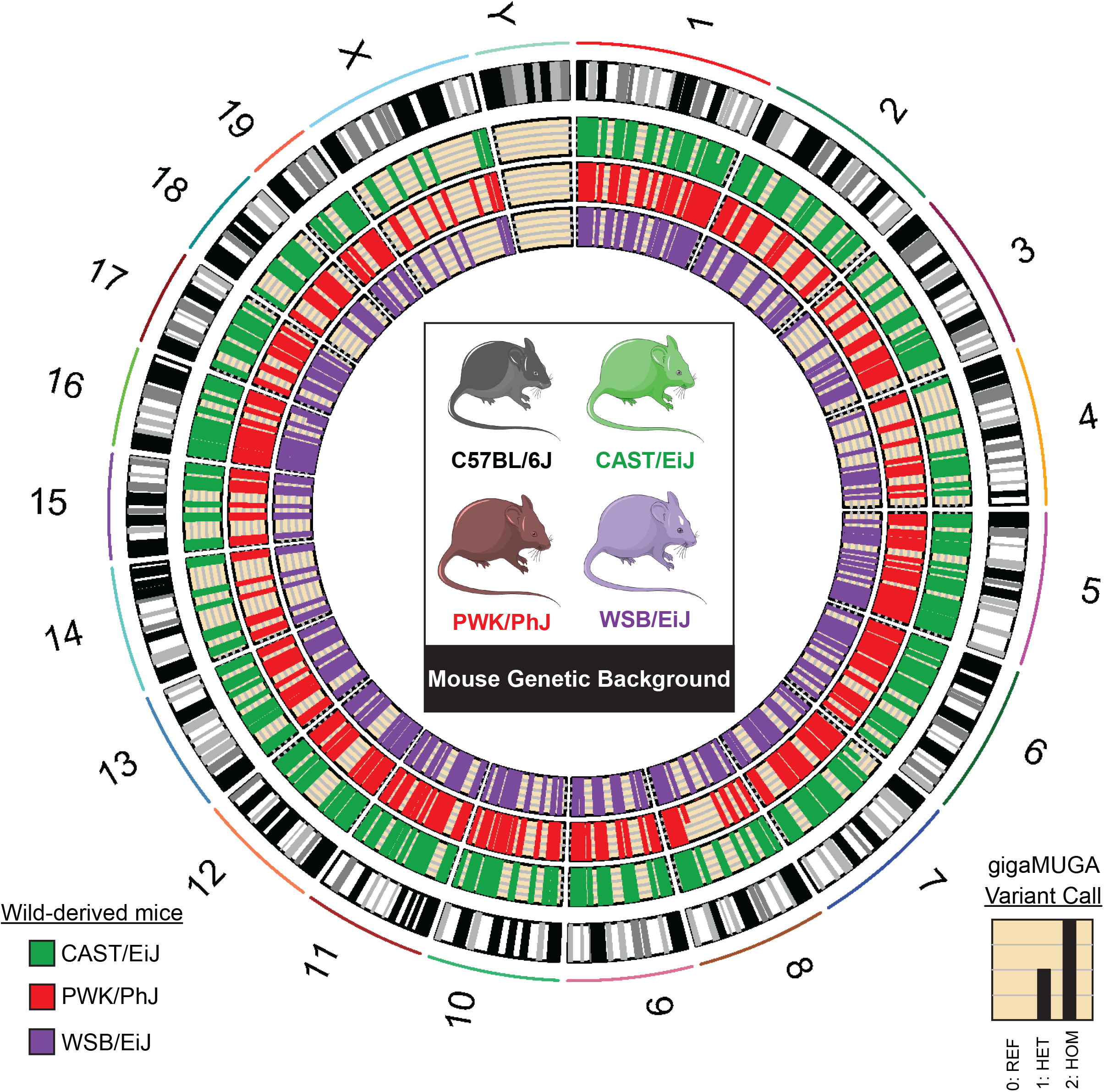
Variants in Wild-derived Genetic Backgrounds within AMP-AD Nominated Target Genes. Classical inbred mouse model C57BL/6J and three wild-derived mouse genetic backgrounds (CAST/EiJ, PWK/PhJ, and WSB/EiJ). Variants in wild-derived mice were called using the Mouse Universal Genotyping Array (gigaMUGA) relative to the reference genome (C57BL/6J) and recoded (0:reference call, 1:heterozygous variant, 2:homozygous variant). Wild-derived mice contain 5,810 variants in the 537 “Nominated Target Genes” from the Accelerating Medicines Partnership Program for Alzheimer’s Disease (AMP-AD) consortium (Accessed March 1, 2021).

In total, we found 5,792 variants in 537 nominated target genes (Supp. File 1A-B). Across all three wild-derived mice in this study, we found genotyped variants in 401 of the total 537 nominated target genes. While a large portion of these variant calls were identical across CAST, PWK, and WSB mice (2,601 out of 5,792), there were a number of strain-specific variants. For example, within Inositol polyphosphate-5-phosphatase D (*Inpp5d*;chr1:87620312-87720507), there were five genotyped variants. One SNP was shared between all three wild-derived mice, three SNPs were shared only by CAST and PWK, and one SNP was heterozygous in CAST and PWK but homozygous in WSB (Supp. File 1C). This demonstrates the genetic heterogeneity of these mouse models within key genes studied in AD.

Cytochrome P450 3A43 (*Cyp3a43*; chr5: 137890932-146113285) contained the most genotyped variants with 563, only 163 of which were shared among all wild-derived mice. 136 of the 537 AMP-AD target genes did not contain any genotyped variants. More information about all variants in these wild-derived mice is available on the Mouse Genome Database (http://www.informatics.jax.org). While our description is limited to those variants genotyped by our selected Illumina Infinium platform, these data suggest that the genetic heterogeneity of the wild-derived mouse genetic backgrounds could modulate many genes of interest for the study of AD and related dementias.

### Pilot Study to Determine Sample Size for Viral Approach

To preserve the effect of genetic background, we selected to model tauopathy with a viral approach. Expressing mutant tau without the need to backcross allows us to test the effect of a “pure genetic background,” without the need to re-genotype each experimental mouse for all variants of interest. We used an AAV-mediated gene expression model, as described before (Carlomagno et al., 2019, Cook et al., 2015, Kim et al., 2008).

To ensure that we would be statistically powered to test the effect of genetic background, we designed a pilot experiment. One litter of B6 and WSB mice was injected with AAV-hTauP301L. At 6 weeks of age, we then evaluated the effects of genetic background on tau seeding activity using the tau seeding assay biosensor assay. Using an effect size of 20% for the FRET+ signal, a power of 0.8, a group number of 4, and *P* < 0.05, we aimed for a final sample size of at least 8 AAV-hTauP301L injected mice per genetic background (Supp. Figure 2A-C).

### Signature A: Core Tau-responsive Signature Across Genetic Backgrounds

To understand the effect that genetic background has on modeling the expression of human tau, we performed bulk RNA sequencing on the cortex of 6-months-old mice injected with either AAV-hTauP301L or AAV-eGFP (Figure 2A; n = 8/background respectively). Principal component analysis (PCA) demonstrates that the largest contribution to the variation in the transcriptome is genetic background (Figure 2B). These data suggest that the genetic variation across genetic backgrounds is the greatest driver of gene expression. To define differentially expressed genes (DEGs) between AAV-hTauP301L and AAV-eGFP injected mice, we used adjusted *P* < 0.05 and a 1.5-fold cutoff for up- or down-regulated genes.

**Figure 2.**
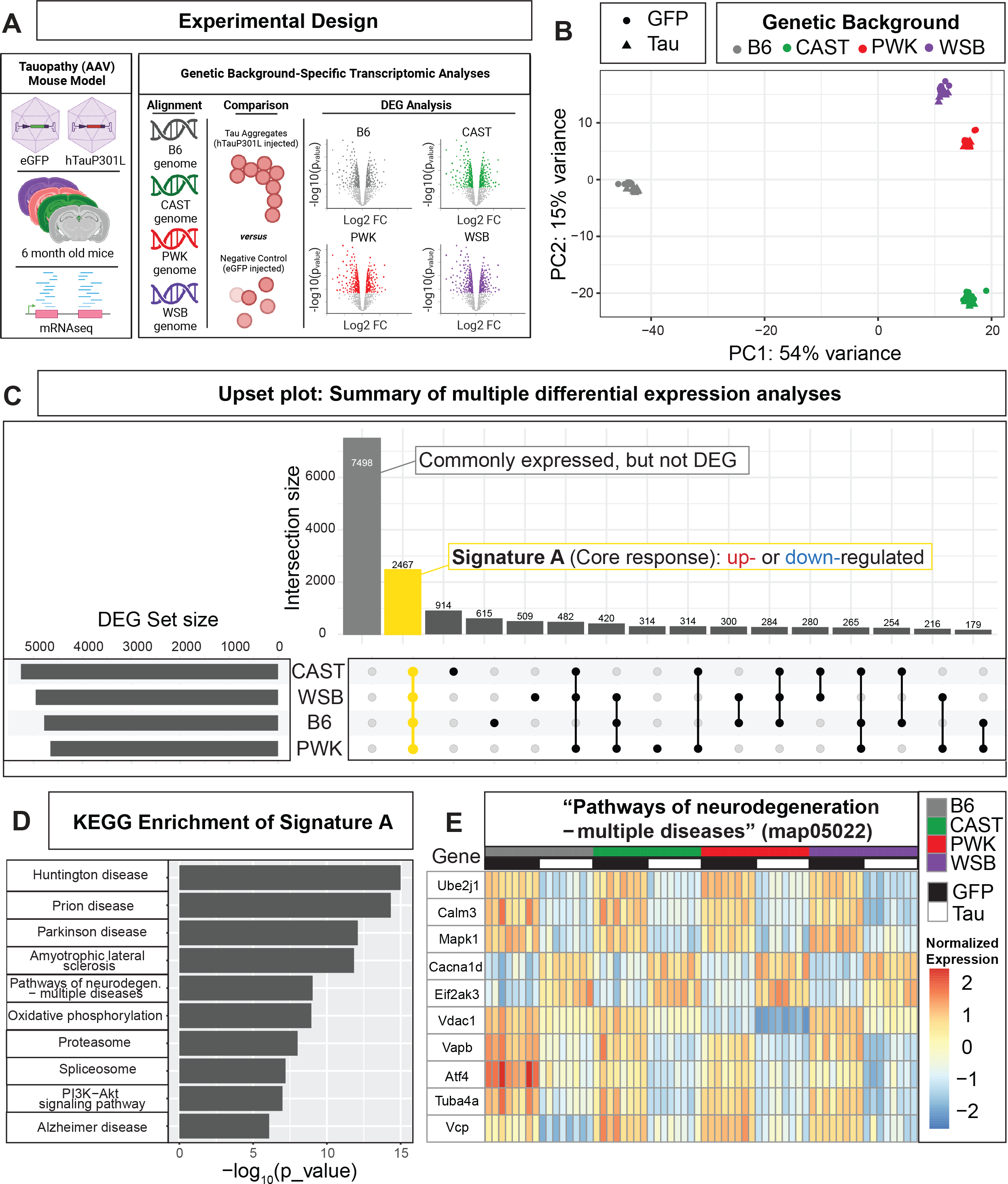
Signature A: Core Tau-responsive Signature Across Genetic Backgrounds. (A) Experimental design to express Tau in B6 and 3 wild-derived mouse strains. In short, AAV-eGFP or AAV-hTauP301L was injected into mice of each genetic background. At 6-months old, brain tissue was collected and analyzed via mRNA-sequencing. Reads were aligned to each strain’s respective genomes. Differential gene expression revealed up-regulated (FC > 1.5, p_adj < 0.05) and down-regulated (FC < − 1.5, p_adj < 0.05) in hTauP301L-injected mice compared to GFP-injected controls (n = 32/AAV injection group). (B) Principal component analysis shows the genetic background drives variation in the transcriptome (n= 8/background/AAV injection group). (C) Upset plot to summarize multiple differential expression analyses: Differential expression (hTauP301L vs eGFP) was performed for each strain (see Supplemental Figure 1, Supplemental File 2A-D). Signature A (highlighted in yellow) was identified as the intersection of differentially expressed genes (DEGs) shared across genetic backgrounds. These 2,467 DEGs were shared across all 4 genetic backgrounds. Other intersections are provided as a resource (Supplemental File 2E). (D) KEGG enrichment of Signature A is significantly enriched for neurodegeneration-related terms and map05022 “Pathways of neurodegeneration” (yellow, DEG intersection: 168/471 genes). See the supplemental information for a summary of all enrichment analyses (Supp. File 2J). (E) Heatmap of the top 10 DEGs in “Pathways of neurodegeneration – multiple diseases” (map05022) shows the conserved response to AAV-hTauP301L injection in Signature A.

Comparisons were made between AAV-hTauP301L and AAV-eGFP injected mice for each genetic background independently (Supp. Figure 1A-D; Supp. File 1A-D). There are a number of genes that are specific to each genetic background (Lilue et al., 2018). Our resource only includes genes that were identified with at least 10 total read counts across all samples of a given genetic background. We identified a total of 4,784 DEGs in B6, 5,260 DEGs in CAST, 4,657 DEGs in PWK, 4,958 DEGs in WSB (DEG Set Size, Figure 2C). Of these DEGs, 2,467 genes were commonly expressed across all genetic backgrounds and were identified as DEGs in all genetic backgrounds (Yellow highlighted Intersection, Figure 2C). The Upset plot also shows genes that were commonly expressed but not DEGs and DEGs shared between different combinations of genetic backgrounds (Figure 2C). Gene sets unique to each background (i.e. CAST-only DEGs, n = 914) are available in a supplemental table (Supp. File 1E). These data suggest a large part (2,467 genes) of what we call the “Signature A: core tau signature” is resistant to the variation between genetic backgrounds.

To understand which genes are part of “Signature A”, we performed enrichment analyses. Kyoto Encyclopedia of Genes and Genomes (KEGG) enrichment analysis of this gene set (2,467 genes) was significantly enriched for several neurodegenerative terms (Figure 2D; Supp. Figure 1E; Supp File 2J). As an example, KEGG term “Pathways of neurodegeneration – multiple diseases” demonstrates a pathway that is conserved across all genetic backgrounds in this study (Figure 2E). A total of 168 genes out of the 471 genes in the pathway (“Pathways of neurodegeneration -multiple diseases” KEGG map05022) follow this genetic background-independent effect (Supp. Figure 1F). For example, *Ube2j1, Calm3*, and *Mapk1* genes were lowly expressed similarly in B6 and all three wild-derived mice injected with AAV-hTauP301L, compared to the AAV-eGFP injected control group (Figure 2E). These data demonstrate that the “core tau signature” includes many targets that are already implicated in neurodegenerative diseases.

### Signature B: Unique Tau Signature in Wild-Derived Genetic Backgrounds

While the presence of DEGs is informative when comparing genetic backgrounds, we were also interested to discover novel targets for tauopathy that may be present only in wild-derived mice. To do this, we performed DEG analysis using the genetic background as a covariate with injection type (~Injection+GeneticBackground+Injection:GeneticBackground). This approach differs from the differential expression analysis to identify Signature A as it identifies DEGs that are not shared across the genetic background in response to tau. A total number of 79 DEGs were identified in CAST (Effect: Tau.CAST; Supp. File 1F), 51 DEGs in PWK (Effect: Tau.PWK; Supp. File 1G), 53 DEGs in WSB (Effect: Tau.WSB; Supp. File 1H). These data suggest that there exist some novel responses to Tau present only in wild-derived mice.

By deciding to calculate DEGs with a genetic background as a covariate, we were able to identify tau response genes that are specific to the wild-derived strains. A Venn diagram of the DEGs shows how many genes are shared across these wild-derived strains (Figure 3A). There were 17 genes shared by CAST, PWK, and WSB (Figure 3B) that were not differentially expressed in B6 mice. As an example, Cilia and flagella associated protein 74 (Cfap74) was up-regulated in wild-derived mice injected with AAV-hTauP301L compared to AAV-GFP injected mice of the same genetic background (Figure 3C). As more risk genes are characterized in the study of tauopathy, it is critical that these genes can be modeled on backgrounds other than B6.

**Figure 3.**
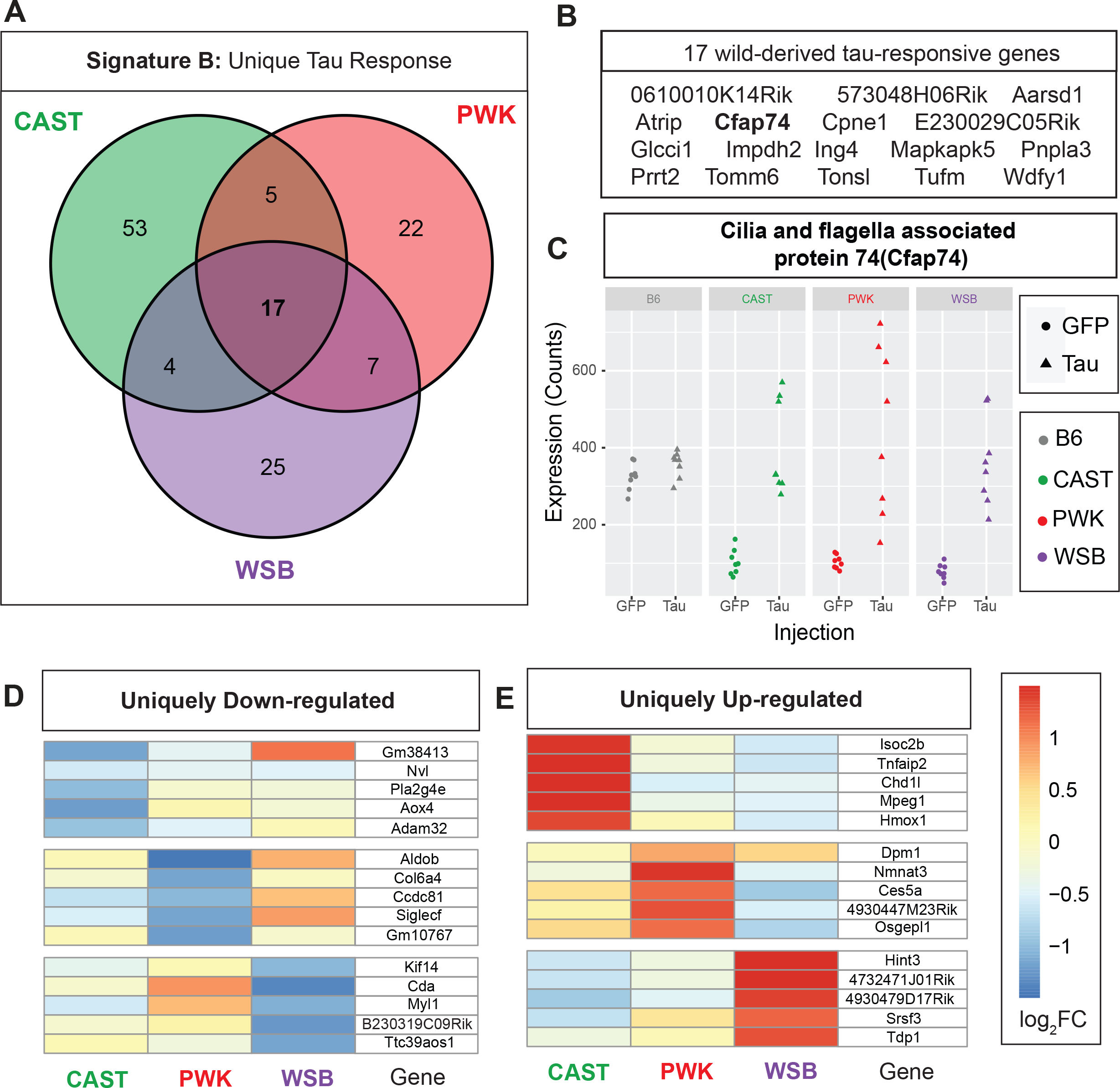
Signature B: Unique Tau-responsive Signatures in Wild-Derived Genetic Backgrounds. Differentially expressed genes (DEGs) specific to wild-derived background (~Injection+GeneticBackground+Injection:GeneticBackground; Benjamini Hochberg adjusted p-value < 0.05, FC > 1.5) were calculated for hTauP301L-injected mice relative to eGFP-injected controls. (A) 133 in total DEGs were identified in one or more wild-derived backgrounds. (B) 17/133 DEGs in Signature B were shared by all three wild-derived backgrounds. (C) Cilia and flagella associated protein 74 (*Cfap74*) and 16 other wild-derived DEGs are not differentially expressed in B6 mice. There are 53 CAST-specific DEGs, 22 PWK-specific DEGs, and 25 WSB-specific DEGs are only differentially expressed in one wild-derived background. The (D) top 5 down-regulated and (E) top 5 up-regulated in each background are shown in a heatmap colored by log2FoldChange between hTauP301L-injected and eGFP-injected mice. See supplemental files for all background-specific DEGs (Supp. File 2F-H).

Uniquely down-regulated (Figure 3D) or up-regulated (Figure 3E) DEGs in just one wild-derived strain are rare phenomena, especially in comparison to Signature A which is comprised of 2,467 DEGs. While there has been no precedent for private DEGs that have a large effect on modeling tauopathy, our data shows some divergent responses to tau. Although the number of wild-derived specific DEGs in Signature B is not large enough to reach significance in enrichment analyses, taken together, these 183 genes in Signature B are enriched for several pathways that have not been studied well in the context of tauopathy (Supp. File 2K).

### Tau Seeding Activity is Modulated by Genetic Background Independent of Tau Expression Level

To investigate whether the pathogenesis of tau aggregates differs across genetic backgrounds, we decided to measure the proteopathic tau seeding activity using an *in vitro* biosensor cell assay. A cell line expressing the repeat domain (RD) of tau conjugated to either a cyan fluorescence protein (CFP) or yellow fluorescent protein (YFP) was transfected with brain lysate from AAV-hTauP301L mice of each genetic background. Fluorescence resonance energy transfer (FRET) signal occurs when tau seeds form due to the proximity of CFP and YFP molecules. FRET+ signal is then measured by fluorescence activated cell sorting (FACS) as a proxy for tau seeding activity (Figure 4A). Sample size was determined based on our pilot experiment (Supp. Figure 2A-C) in order to design the current study (Supp Figure 2D).

**Figure 4.**
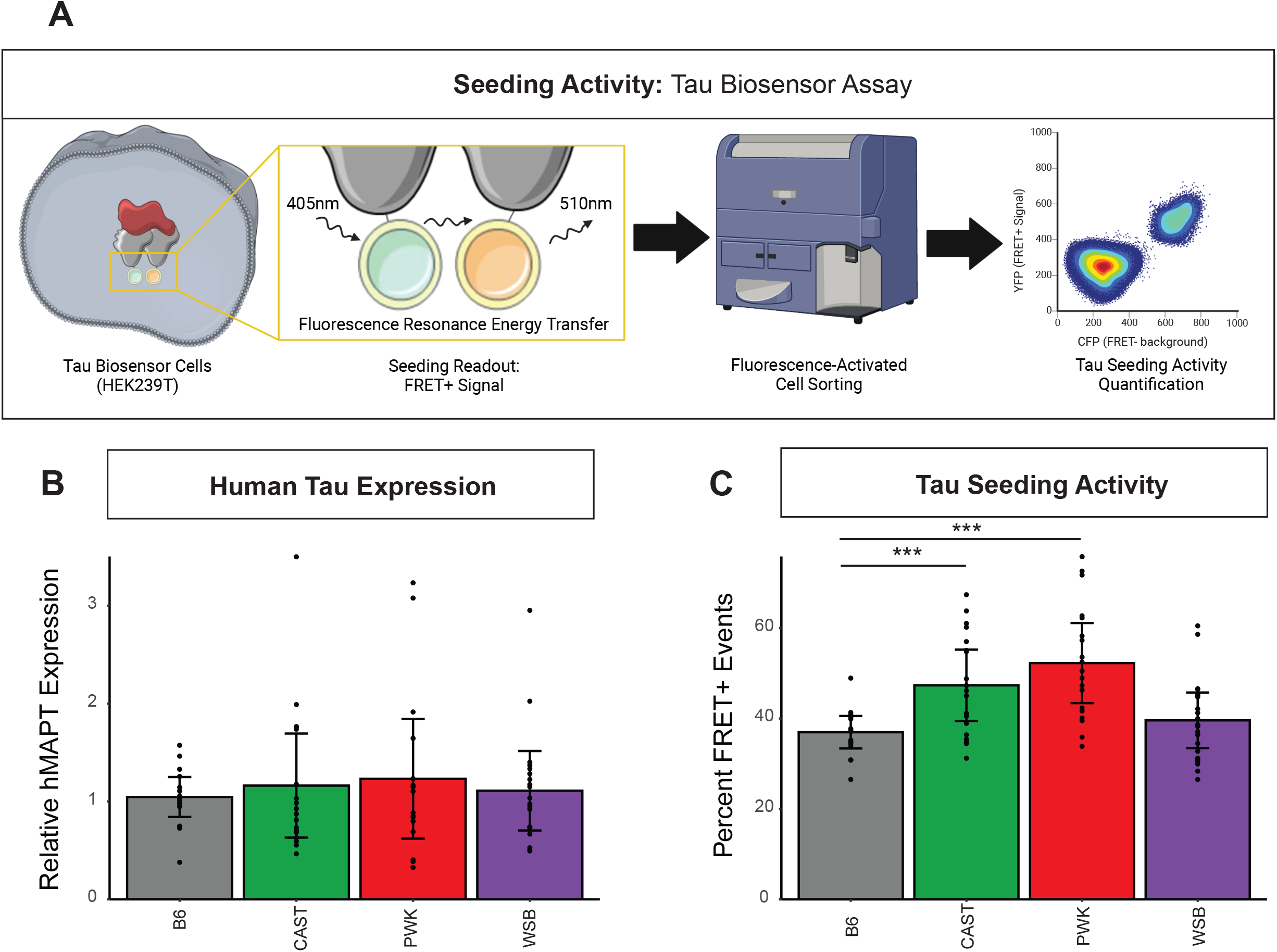
Tau Seeding Activity is Modulated by Genetic Background Independent of Tau Expression Level. (A) Paradigm of Tau biosensor cells to measure seeding activity assay. HEK-293T cells containing CFP- or YFP-conjugated tau are transfected with brain lysate from hTauP301L-injected mice for 24 hours. Biosensor cells are then collected and FRET+ signal is measured via Fluorescence-Activated Cell Sorting (FACS) as a proxy for tau seeding activity. (B) Human tau expression was measured using quantitative PCR. Relative hMAPT expression was calculated relative to GAPDH and showed no effect of genetic background (n = 69, F_3,66_ = 0.234, p = 0.87). (C) Tau seeding activity was measured as the % of HEK-293T biosensor cells in the final FRET+ (YFP) gate using FACS. Tau seeding activity was a significantly effected by genetic backgrounds (n = 81, F_3,78_ = 9.237, p = 2.67 x10-5). Tukey HSD post-hoc test revealed elevated tau seeding activity in CAST and PWK relative to B6 (***: *p* < 0.001).

To ensure consistency in viral expression of the hTauP301L, we measured human *MAPT* expression from the cortex of B6, CAST, PWK, and WSB mice. We found no significant effect of genetic background on the expression of Tau by our AAV construct (Figure 4B; F_3,66_ = 0.234, *p* = 0.87). Qualitative analysis of regional human tau expression (Supp. Figure 2E), the presence of AT180+ (pThr231/pSer235) aggregates (Supp. Figure 2F), and specific HT7+ bands (Supp. Figure 2G) suggest no effect of genetic background. Genetic background did not affect the levels of total tau and pTau231(Supp. Figure H). These data suggest that the genetic variations across different mouse genetic backgrounds do not influence our ability to express tau using intracerebral ventricle AAV injection.

Interestingly, genetic background significantly affected tau seeding activity when we used cortical tissue lysates as seeding agents. Percent FRET+ events measured by flow cytometry were modulated by genetic background (Figure 4C; F_3,78_ = 9.237, *p* = 2.67 × 10^−5^). Tukey Honest Significant Difference (HSD) post-hoc testing revealed a significant increase in PWK and CAST mice relative to AAV-hTauP301L-injected B6 controls (*p* < 0.001). However, there was no significant difference between B6 and WSB AAV-hTauP301L-injected mice (*p* > 0.05). Unlike tau seeding activity from cortical lysates, no effect of genetic background on tau seeding activity was observed with hippocampal tissue lysates (Supp. Figure 2I). To ensure the rigor of our analysis, we replicated our tau seeding assay data. The replication of tau seeding activity assay showed a high correlation between technical replicates in the cortex (R^2^ = 0.7988) and hippocampus (R^2^ = 0.8701; Supp Figure 2J). Taken together, our data suggest that the genetic heterogeneity across wild-derived mice exacerbates the prion-like action of tau, specifically in the brain cortex.

### Signature C: Tau Seeding-Associated Signature in PWK and CAST strains

The finding that tau seeding activity varies across genetic backgrounds compels us to ask whether there is an associated tau seeding signature. We performed Weighted Gene Co-expression Network Analysis (WGCNA) to investigate which genes may be correlated with this difference in seeding activity. Using our FRET data as a trait for “module-trait” relationship analyses, we aimed to elucidate which genes may be involved in tau seeding. We identified 60 modules of co-expressed genes (Supp. File 1I; Supp. Figure 3A-C) and tested their correlation to traits including: whether mice were injected with AAV-eGFP or AAV-hTauP301L (Injection), whether mice were B6 or one of the wild-derived backgrounds (wild-derived), biological sex (Sex), and seeding activity (FRET). Of these 60 modules, 11 were significantly associated with seeding activity (Supp. Figure 3D, *p* < 0.05). Importantly, none of the 11 modules demonstrated a significant effect of sex.

Each of these 11 modules with a significant Module:TraitFRET relationship may contain genes that explain the increase in seeding activity seen in PWK and CAST mice. We focused on those that were most highly associated via Pearson Correlation (Figure 5A, R^2^: Positively Correlated 0-1, Negatively Correlated -1-0). Although the cyan module was most positively correlated to the FRET measurement (Figure 5A, R^2^ = 0.65, *p* = 2 × 10-8), it showed no effect of wild-derived background (R^2^ = 0.0097, *p* = 0.9). The darkorange module is the second most postively correlated to FRET measurement (Figure 5B, R^2^ = 0.54, *p* = 7 × 10^−6^) and showed a significant effect of wild-derived background (R^2^ = 0.27, *p* = 0.03). Eigenvalue of the genes in the both modules demonstrates that the effect of wild-derived background was not present in the cyan module (Figure 5A), but did appear in the darkorange module (Figure 5B). For this reason, we focused on the darkorange module to describe the effect of genetic background on tau seeding activity “Signature C.”

**Figure 5.**
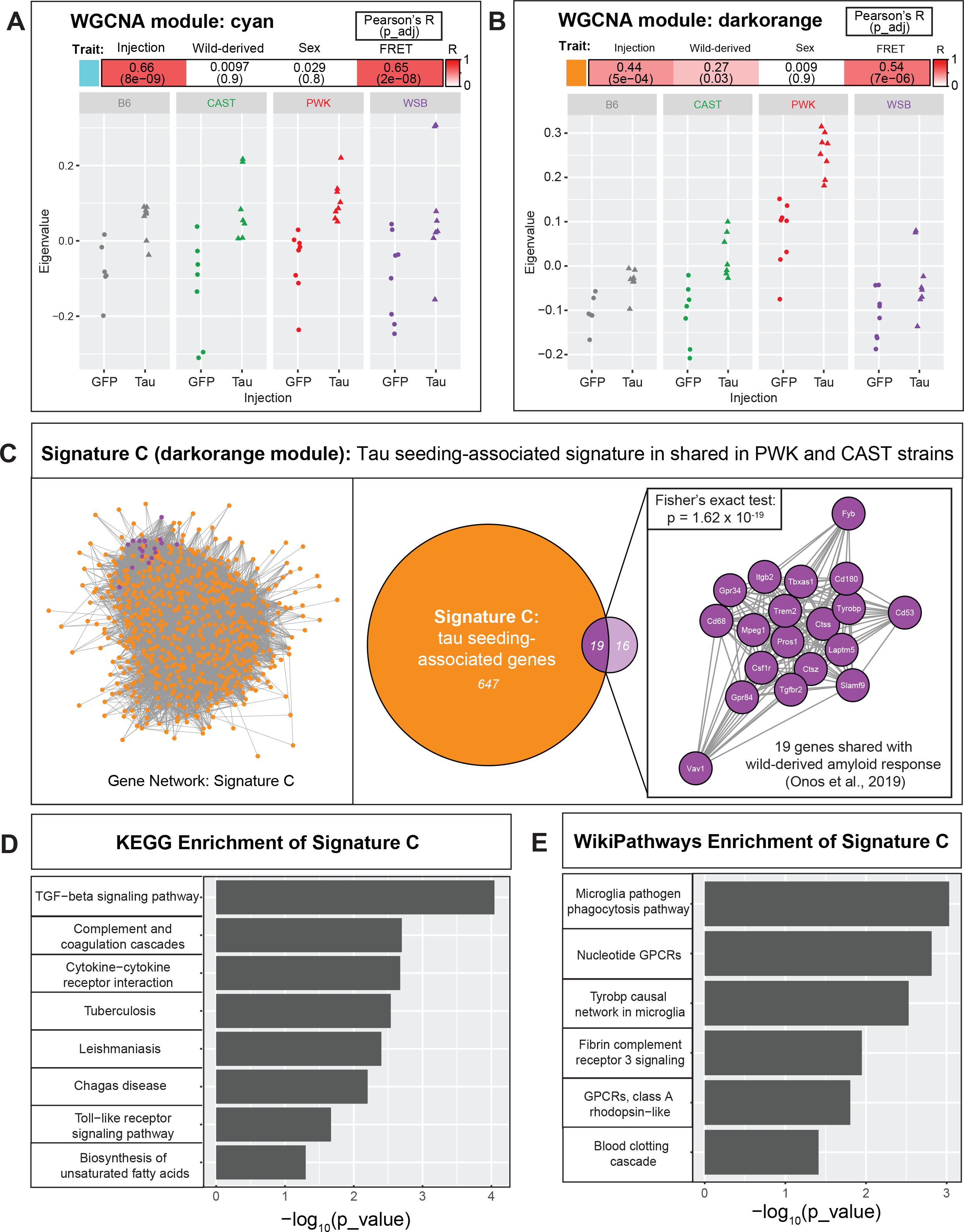
Signature C: Tau Seeding-Associated Signature in PWK and CAST strains. Gene module detection was performed using Weighted Gene Co-expression Network Analysis (WGCNA) from hTauP301L-injected and eGFP-injected mice. Module-trait detection revealed the top two tau seeding-associated modules (A) cyan and (B) darkorange. Discovery of associated modules were prioritized based on association with injection type (Injection, 1^st^ column), genetic background (Wild-derived, 2^nd^ column), sex (Sex, 3^rd^ column), and tau seeding activity (FRET, 4^th^ column). Statistics for module-trait association include Pearson’s R and Benjamini-Hochberg adjusted p value. A full list of all 60 modules and their module-trait correlation can be found in Supplemental Figure 3.(C) Darkorange module was renamed “Signature C”: Gene network represented as nodes with edge distance representing topological overlap matrix (TOM) score from WGCNA. Neurodegenerative hub-genes (purple) were detected by comparing Signature C (darkorange) and wild-derived amyloid response (Fisher’s exact test: p = 1.62 x 10^−19^; Onos et al., 2019). (D) KEGG enrichment and (E) WikiPathways enrichment of Signature C reveal microglia-related terms. See supplemental information for WGCNA parameters (n = 60 samples after outlier detection).

The darkorange module contains a total of 666 genes that we found to be associated with the effect of genetic background on tau seeding activity (Figure 5C, highlighted in orange). A similar effect of the genetic background was found via WGCNA when modeling Aβ (Onos et al., 2019). In comparison to their 35-gene module, we found that 19 of their genes were significantly enriched in our darkorange module (Figure 5C, highlighted in purple; Fisher exact test, p = 1.62×10^−19^). Interestingly, these genes that are affected by genetic background in both amyloid and tau studies include key microglial genes (*Trem2*, *Tyrobp*, *Tgfbr2*) and the complement cascade genes (*C1qa*, *C1qb*, *Pros1*) (Figure 5C). This finding suggests that both pathways are sensitive to genetic context when modeling both hallmarks of AD.

We further characterize this tau seeding-associated signature using enrichment analysis. Enrichment analysis of these genes revealed KEGG terms associated with immune response (Figure 5E) and WikiPathways associated with microglia (Figure 5F). These data suggest that the immune system, specifically microglia, may be implicated in the increase in seed-competent tau observed in CAST and PWK mice. Additional enrichment analyses of Signature A, Signature B, and Signature C can be found in the supplemental information (Supp. File 2J-L).

## DISCUSSION

Recent studies in AD have shown that mouse genetic background can modulate Aβ accumulation (Onos et al., 2019), immune response (Yang et al., 2021), and tau propagation (Dujardin et al., 2022). To better understand how genetic background influences tauopathy, we aimed to create a resource of core- and unique-transcriptional signatures to tau expression based on mouse genetic background. Additionally, we were interested in determining if the seeding activity of tau was modified by genetic background. To better understand the pathogenicity of Tau aggregates, it is important to investigate the initial seeding of tau and subsequent spreading/propagation, similar to studying Aβ in AD and α-synuclein in Parkinson’s disease (Peng et al., 2020). We report that the cortex of wild-derived CAST and PWK mice has significantly higher prion-like proteopathic seeding activity of tau compared to that of B6 controls. To better understand the mechanisms involved, we performed a network analysis that implicated microglia in this strain-specific seeding activity. Our data suggest that mouse genetic background is an important factor when studying immune responses to pathological tau species.

For this study, we selected three wild-derived genetic backgrounds (CAST, PWK, and WSB) to compare to B6. Using a genotyping array, gigaMUGA, we report that these three wild-derived strains contain many variants in the nominated targets from the AMP-AD database (Figure 1). A comprehensive list of all variants in these wild-derived mice and 85 other strains can be found in the Jackson Laboratory’s Mouse Genome Database (Blake et al., 2021). Transcriptomic data from Onos and colleagues showed a compelling effect of these three wild-derived strains on amyloid accumulation (Onos et al., 2019, Yang et al., 2021). To compare the response to the two hallmarks of AD, amyloid and tau, we decided to investigate these same mouse strains. Our data suggest that these wild-derived strains are an ideal resource for investigating the contribution of genetic variation to the study of AD and other tauopathies.

With AD mouse models, transcriptional signatures have been an important experimental readout. Several studies have shown that this hypothesis-generating, unbiased readout can be used to investigate Aβ accumulation (Sierksma et al., 2020), region-specific expression of tau (Castanho et al., 2020), and activated immune response (Kang et al., 2018). Previously, genetic background has been shown to influence the amount of tau and the presence of at least one phospho-epitope in tau transgenic models (Eskandari-Sedighi et al., 2017, Yanagisawa et al., 2021, Bailey et al., 2014). However, no transcriptional information or mechanism of action has ever been proposed. To test the effect of wild-derived genetic background and generate hypotheses about the responsible mechanisms, we used transcriptional signatures as our main readout. Importantly, our study shows that the core transcriptional response to tau across different genetic backgrounds is enriched for pathways of neurodegeneration (Figure 2). This finding suggests that the fundamental pathways involved in studying mouse models of dementia do not change across genetic backgrounds. As preclinical studies continue to make direct comparisons between human and mouse transcriptomics (Monzon-Sandoval et al., 2022, Onos et al., 2022), our list of core genes can be interpreted as robust tau-responsive genes for future study.

However, we argue that it is just as important to understand what transcriptomic response is modulated by different genetic backgrounds. Previous studies of Trem2 on mixed background mice indicated an allele inherited by the SJL strain that unknowingly introduced a missense mutation (Yang et al., 2021). In previous studies, we have addressed this issue by excluding mice that are homozygous for the SJL allele from analysis (Karahan et al., 2021). However, for investigators planning to study novel risk genes, it would be impossible to control for every naturally occurring variant. We propose using these differences, specifically in the study of tauopathy, to our advantage. Should researchers consider focusing on any gene of interest, it is critical that we first understand what aspects of mouse biology are causing the genetic background to modulate tau phenotypes. We report a unique or “segregating” response to tau in wild-derived mice (Figure 3). For example, WSB mice appear to have a unique down-regulation in two genes involved in motor transport, *Kif14* and *Myl1* (Figure 3D). For those studying the role of motor transport in AD (Gan et al., 2020), the WSB background may provide insights that would not be observed using B6 mice. Other conclusions from the wild-derived unique responses could explain unexpected negative data when using B6 mice. This would be one of the barriers to translating research findings into humans

Lastly, to understand if genetic background modulates the pathogenicity of tau, we investigated the effect of mouse strain difference on the tau seeding activity. We found an increase in tau seeding activity in the cortex of CAST and PWK mice (Figure 4). More research is necessary to understand why this increase in seeding activity is occurring and why we observed the effect of genetic background only in the cortex and not in the hippocampus. It is possible that modifiers or interactors of tau exist in CAST and PWK mice. Our previous research has demonstrated that interactors like Bassoon (*Bsn*) contribute to the ability of tau to seed and induce neurotoxicity (Martinez et al., 2022). As a pathological readout, seeding activity has been shown to identify the action of high molecular weight tau (Martinez et al., 2022) and even differentiate between specific conformers in 3R/4R tau diseases (Kraus et al., 2019). Importantly, our network analysis identified a module of genes associated with this increase in CAST and PWK mice (Figure 5). Our enrichment analysis suggests the importance of microglia (Figure 5D, E). However, this finding is based on bulk transcriptomic data from wild-derived mouse strains. An increasing amount of work is currently going into identifying disease- and context-specific glial states (Keren-Shaul et al., 2017, Paolicelli et al., 2022, Ezerskiy et al., 2022). Single cell RNA sequencing will be necessary to better understand which microglia cell types are involved in this phenotype. Interestingly, we see a considerable overlap when comparing the tau-seeding associated genes in wild-derived mouse strains to a previous study of amyloid response in wild-derived strains (Onos et al., 2019). These include *Trem2*, *Tyrobp*, *Tgfbr2*, and *Cd68*.

In conclusion, we have described strain-specific variants identified via Illumina Infinium platform and three transcriptional signatures identified via RNA sequencing. First, a core tau-responsive signature that is not affected by genetic background (Signature A). Second, a unique response to tau that may indicate wild-derived mice should be used to study specific risk genes (Signature B). Third, a tau seeding activity associated signature that implicates microglia (Signature C). Our data provide a resource for investigating tau in mouse models of AD and other tauopathies (Figure 6). Given that most therapeutic approaches are tested in mice before progressing to clinical trials, including wild-derived mice may enhance the translatability to treating patients with different genetic backgrounds.

**Figure 6.**
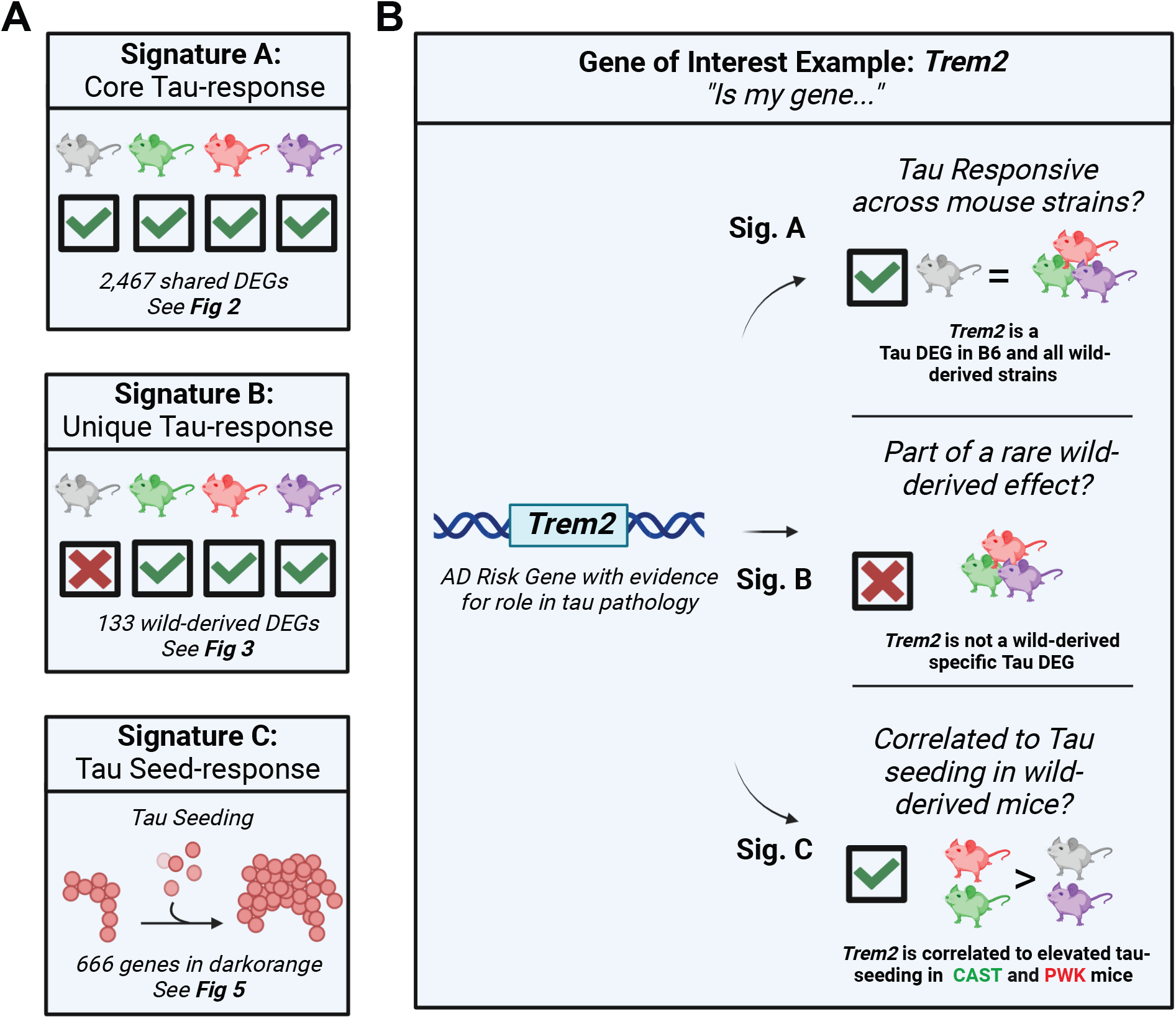
Resource: Guideline to Select a Mouse Genetic Background to Study Tau. (A) Signatures A-C of this study represent a core response to expressing AAV-hTauP301L (Signature A), wild-derived specific response to AAV-hTauP301L expression (Signature B), and a tau seeding-associated module (Signature C). (B) Given a gene of interest, the resources in this paper can guide genetic background selection for functional studies in mice. For example, *Trem2*, a gene with strong evidence for a role in tau pathology, is present in Signature A (Core Tau-response) and Signature C (Tau seed-response). Based on this evidence, while *Trem2* is differentially expressed in response to tau across all mouse strains, there is a possibility that it is involved in a CAST- or PWK-specific reaction to tau seeds.

## MATERIALS AND METHODS

### Mouse strains and Genotyping

This study was designed to investigate the role of genetic background in the pathogenesis of tauopathy. To achieve this, we purchased breeders from genetically diverse mouse strains from the Jackson laboratory (C57BL6/J: Stock #000664 ,CAST/EiJ: Stock #000928 ,PWK/PhJ: Stock #003715, WSB/EiJ: Stock #001145). All procedures and animal work were approved by the Indiana University School of Medicine Institutional Animal Care and Use Committee (Protocol 21149).

Tail samples from a pilot cohort were collected and sent to GeneSeek (Neogen) for genotyping on the gigaMUGA (Mouse Universal Genotyping Array) platform. This array contains 143,259 SNP and CNV markers that were selected to be informative in wild-derived mice and multiple *Mus* species (Morgan et al., 2015). The results published in this study are in part based on data obtained from Agora (https://agora.adknowledgeportal.org/), a platform initially developed by the NIA-funded AMP-AD consortium that shares evidence in support of AD target discovery. In argyle (Morgan, 2015), variants were recoded and filtered based on the position of the AMP-AD nominated target genes (Accessed March 1, 2021). Variants were then visualized using RCircos (v1.2.2).

### Intracerebroventricular Injections of Adeno-associated Virus

To model tau aggregation, we injected mice of each genetic background with either AAV-hTauP301L (AAV9-CBA/CMV-hTauP301L-WPRE-polyA) or AAV-eGFP (AAV9-CBA/CMV-eGFP-WPRE-polyA). Sample size was determined by the pilot study described in the results section. Final sample size reached our requirement based on power analysis (B6 = 20, CAST = 20, PWK = 19, WSB = 24). We selected AAV9 which has been shown to have high intracellular expression without an effect of mouse genetic background (He et al., 2019).

A full protocol for this approach was previously published (Kim et al., 2014, Passini et al., 2003). In brief, breeder cages were checked three times daily to ensure injection occurred between 12-24 hours after birth (Postnatal Day 0, P0). P0 mice were cryo-anesthetized for 8 minutes on ice. Using a 32-guage needle, 2 millimeters deep injections were made into each lateral ventricle (0.8-1mm later from the sagittal suture and hallway between lambda and bregma) at a 45-degree angle. A total of 2 μL of virus were injected per ventricle (4 x 10^10^ viral particles/mouse) to express each construct. The injection was performed slowly, and the needle is held in place for an additional 30 seconds. Upon removal of the needle, if more than 0.2 μL of the virus leaks out of the injection site, the animal is immediately euthanized. Surviving mice were placed on a warming pad until the pup begins to move and were promptly returned to their parent cage.

### Tissue harvesting and sample preparation

At six months of age (182.1 +/− 4.9 days, mean +/− standard deviation), mice were anesthetized using carbon dioxide for 2.5 minutes. Brains were promptly removed, and the left hemisphere was fixed in 4% paraformaldehyde for 24 hours at 4°C to be stored for histology. Tissue samples were embedded in paraffin and sectioned at Histology Lab Service Core at the Indiana Center for Musculoskeletal Health. Five-micrometer-thick coronal sections (at Bregma −1.46, −1.94, and −2.46 mm) were transferred to charged microscope slides and stored at room temperature. The Anterior Cortex, Posterior Cortex, Hippocampus, and Cerebellum were dissected from the right hemisphere and flash-frozen with liquid nitrogen. Samples were stored at −80°C.

### Protein preparation

Samples were weighed and prepared in 1X Tris-buffered saline (TBS) at 100 mg of tissue per milliliter of lysate. After a brief gentle mechanical dissociation, samples were aliquoted for either RNA or protein extraction. The aliquot designated for protein extraction was homogenized via sonication and centrifuged at maximum speed for 15 minutes at 4C. The supernatant, referred to as “TBS-soluble,” was then normalized to 2.0 mg/mL via Bicinchoninic Acid (BCA) assay (Thermo Scientific) and stored at −80C until analyzed.

### Tau-seeding assay

The seeding assay was performed using TauRD P301S FRET Biosensor Cells (Holmes et al., 2014). In brief, we obtained HEK293-T cells expressing truncated TauP301S containing only the Repeat Domain (RD) fused to either Cyan- or Yellow-Fluorescent Protein (CFP, RYP) (ATCC CRL-3275). These biosensor cells were plated in a 96-well plate at 30,000 cells per well and incubated at 37°C overnight. After 24 hours, cells were transfected with 20 μg of TBS-soluble brain lysate using Lipofectamine 2000. After an additional 48 hours at 37°C, cells were harvested and FRET+ signal was measured via Flow Cytometry (BD LSRFortessa X-20 with High Throughput Sampler). Data analysis was performed in FlowJo (v10.0). Our gating strategy for singlet selection, CFP background removal, and FRET+ signal (BV510 channel) was performed as previously described (Martinez et al., 2022). Tau-seeding activity was quantified as the percent of total cells with FRET+ signal.

### Western blotting

A total of 15 μg of protein were loaded onto a 4 to 20% TGX gel (Bio-Rad), separated by gel electrophoresis, and transferred onto polyvinylidene difluoride membranes. Membranes were blocked with 5% Bovine Serum Albumin (BSA) in tris-buffered saline (TBS) containing 0.05% Tween20. Blots were probed with mouse anti-human TAU (HT7; 1:50,000; Invitrogen MN1000), chicken anti-GFP (1:1,000; Abcam ab 13970), and rabbit anti-GAPDH (1:10,000; Santa Crus sc-25778) antibodies overnight at 4°C. Membranes were washed with TBS containing 0.05% Tween20 and incubated with anti-mouse, anti-rabbit, or anti-chicken HRP-linked IgG antibodies based on primary antibody host (1:1,000). Membranes were developed via chemiluminescence [ECL Select (GE Healthcare)].

### Tau Protein Quantification

Quantification of total tau protein and phosphor-tauThr231 were measured using a kit from Meso Scale Diagnostics (K15121D). To read both total and phosphorylated tau simultaneously, normalized protein lysate from brain (2.0 mg/mL) was diluted 1:2,000. Signal detection was performed via MESO QuickPlex SQ 120MM and analysis was done using Methodical Mind software (Meso Scale Diagnostics, Rockville, Maryland).

### Histology and immunohistochemistry

Slides containing mounted coronal sections were deparaffinized using xylene. Antigen retrieval was performed with Low pH IHC Antigen Retrieval Solution (Invitrogen) in a pressure cooker set to 100°C for 10 minutes. For 3,3’-diaminobenzidine (DAB) staining, endogenous peroxidation was quenched by incubating slides in a solution containing 10% methanol, 3% hydrogen peroxide in phosphate-buffed saline (PBS) for 10 minutes. Slides were blocked with 5% Normal Goat Serum (NGS) in PBS containing 0.25% Triton X-100. Sections were incubated with primary antibodies overnight at 4°C. Human tau-specific (HT7, 1:2,000; Invitrogen MN1000) and Tau^pThr231/pSer235^(AT180, 1:2,000, Invitrogen MN1040) antibodies were diluted in 2.5% NGS in PBS containing 0.4% Triton X-100. Sections were washed, briefly incubated in 1% BSA, and incubated with biotinylated goat anti-mouse secondary antibody (1:400, Thermo Fisher Scientific) at room temperature for 1 hour. Antibody detection for DAB development was done using the Vectastain ABC Elite (Vector Laboratories, PK6100) and DAB peroxidase substrate kits (Vector Laboratories, SK-4100). Sections were dehydrated and cleared with Ethanol and Xylene and immediately coverslipped with mounting medium containing 20% Xylene (Permount, Thermo Fisher Scientific). Representative images were obtained using brightfield microscopy at the magnifications noted in figure legends (4X, 20X).

### RNA preparation for qPCR and mRNA-seq

Total RNA was extracted from posterior cortex brain tissue using TRIzol (MRC). RNA concentration and quality were determined via Nanodrop 200 Spectrophotometer.

For real-time quantitative Polymerase Chain Reaction (qPCR), cDNA was prepared using a High Capacity cDNA Reverse Transcription kit (Applied Biosystems). qPCR was performed in QuantStudio 3 using the recommended protocol for FAST SYBR (Applied Biosystems) with the following primers: human specific *MAPT* forward TTGCTCAGGTCAACTGGTTT, human specific *MAPT* reverse ACTGAGAACCTGAAGCACCA, mouse *Gapdh* forward AAGGTGAAGGTCGGAGTCAAC, mouse *Gapdh* reverse GGGGTCATTGATGGCAACAATA. Relative mRNA levels were calculated by comparative cycle threshold (ΔΔCt).

For mRNA-seq, total RNA was concentrated and purified using RNA Clean-Up & Concentrator-5 kit (Zymo Research). RNA Integrity Number (RIN) and concentration were determined via TapeStation RNA tape (Agilent). Sequencing was performed by the Center for Medical Genomics at the Indiana University School of Medicine (Indianapolis, IN). Libraries created from 100ng of total RNA using mRNA HyperPrep kit (KAPA). Libraries were then checked for quality and loaded at a concentration of 300 pM on a flow cell for 100 bp paired-end sequencing (S4_200cycle flow cell v1.5). Sequencing was then performed on an Illumina NovaSeq 6000 at an average sequencing depth of ~30 million reads per sample.

### Transcriptomic analyses

Reads were mapped to the respective reference genome of each genetic background (B6-UCSC/refGene mm10, CAST-GCA_001624445.1, PWK-GCA_001624775.1, WSB-GCA_001624835.1) using RNA-seq aligner STAR (v.2.7.10a). See Supplement information for sequencing and mapping statistics (Supp. File 2M-Q). Reads were assigned to genomic features using featureCounts (Liao et al., 2014). Raw read counts were analyzed for either differential expression analysis in DESeq2 (Love et al., 2014) (v1.36.0) or network analysis using Weighted Gene Co-expression Network Analysis (Langfelder and Horvath, 2008) (v1.71).

For differential expression analysis, two separate strategies were applied. To identify a core transcriptional response to expressing hTauP301L, analysis was first done on each genetic background separately. This allowed for strain-specific genes that are not annotated in the mm10 reference genome to be included in our initial analyses. Within each genetic background, genes with a raw read count of less than 10 were filtered out. Differential gene expression was then calculated for AAV-hTauP301L injected mice relative to AAV-eGFP control (B6 = 17,963, CAST = 22,426, PWK = 22,100, WSB = 22,399 genes after filtering). Up- and down-regulated genes were defined using a significance cutoff of 0.05 (Benjamini Hochberg adjusted p-values) and a 1.5-fold change (after *apeglm* effect size shrinkage (Zhu et al., 2019)).

The second differential expression analysis aimed to find unique responses to hTauP301L without the effect of genetic background or tau expression alone. To do this, raw read counts across genetic backgrounds were merged keeping only genes that were annotated in the mouse reference genome (mm10). After merging, 19,468 genes were identified at least once in each genetic background and 17,240 had at least 10 read counts across all genetic backgrounds. These 17,240 genes were used in all downstream analyses (PCA, unique transcriptional response, WGCNA). Note that there were several genes mapped to wild-derived backgrounds that were removed from the analysis because they correspond to more than one gene on the reference genome (CAST = 12, PWK = 12,WSB = 14 multi-mapped genes removed). These were all either predicted genes (“Gm” prefix) with the exception of one small nucleolar RNA (Snora43).

Raw reads underwent variance stabilization transformation (vst) and principle component analysis (PCA) was used to identify potential outliers. Differential gene expression was then performed with genetic background as an interaction (~Injection+GeneticBackground+Injection:GeneticBackground). The goal of this calculation is to identify tau-responsive genes that were dependent solely on genetic background. Up- and down-regulated genes were defined using a significance cutoff of 0.05 (Benjamini Hochberg adjusted p-values) and a 1.5-fold change (after *apeglm* effect size shrinkage) for each interaction term (Tx_Tau_CAST, Tx_Tau_PWK, and Tx_Tau_WSB) with the B6 as a baseline.

We then performed WGCNA to identify modules of co-expressed genes that could explain the variation we reported in tau seeding activity across genetic backgrounds. For this analysis, we returned to the raw count matrices without any normalization or filtering as recommended by the authors of the pipeline (https://horvath.genetics.ucla.edu/html/CoexpressionNetwork/Rpackages/WGCNA/faq.html). After outlier removal in WGCNA, we were left with a total of 60 samples (Supp. Figure 3A). Our data did not reach the suggested scale free topology model fit cutoff of 0.9 (Supp. Figure 3B). A soft power threshold of 6 was selected based on the suggestions by the authors of the pipeline for a dataset with more than 40 samples. A total of 60 modules were identified (Supp. Figure 3C) and 11 of them were significantly associated with FRET+ seeding activity (Supp. Figure 3D; Pearson’s correlation, *p* < 0.05).

Enrichment analyses for core tau signature (Supp. File 3J), unique tau signature (Supp. File 3K), and WGCNA modules (Supp. File 3L) were performed in gProfiler2 (R Client, v0.2.1). Output includes enrichment for Gene Ontology (GO) terms, Reactome (REAC), TRANSFAC (TF), miRTarBase (MIRNA), Human Protein Atlas (HPA), Comprehensive Resource of Mammalian Protein Complexes (CORUM), Human Phenotype Ontology (HP), and WikiPathways (WP).

For enrichment of our signatures against other published datasets, we performed a Fisher’s exact test using the stats package in R (v4.2.1). Given the size of test signature (A), the size of the signature from literature (B), the size of overlap between signature A/B (t), background (n = whole transcriptome), enrichment was considered significant if p < 0.05. The command stats∷dhyper(t:B,A,n-A,B) returned the p-value for Fisher’s exact enrichment.

### Statistical analysis and Figure Creation

For analysis of tau pathology, analysis was done via one-way Analysis of Variance (ANOVA) followed by Tukey Honest Significant Difference (HSD) post-hoc test. Statistical tests are reported in the figure legends with sample size, F statistic, degrees of freedom, and *p* value. Where appropriate, figures are labeled with the exact *p* value (*p* > 0.05), * (*p* < 0.05), ** (*p* < 0.01), *** (*p* < 0.001). All analysis was done in R (v4.2.1) and figures were created using ggplot2 (v3.3.6),Cytoscape (v3.9.1), and BioRender.com.

### Data availability

Data files from mRNAseq analysis will be made publically available on Gene Expression Ombinus (GEO) upon publication. All remaining data can be found in the supplemental information in this manuscript or can be made available upon request.

### Supplemental data

Supplemental figures include a summary of genetic background-specific transcriptomic analyses not shown in the main figures (Supp Figure 1), a description of a pilot study to determine sample size (Supp. Figure 2A-C), characterization of our AAV-injected tau module suggesting no effect of genetic background on the expression of human tau or tau seeding activity in the hippocampus (Supp. Figure 2D-J), and a summary of WGCNA analysis (Supp. Figure 3). Supplemental files contain wild-derived AMP-AD variant information (Supp. File 1) and summaries of transcriptomic analyses (Supp. File 2).

## ACKNOWLEDGEMENTS

This work was supported by National Institute of Health Grants R01 AG077829, R01 AG071281, RF1 AG074543, R21 AG072738 (J.K.); NIH Predoctoral Fellowship provided through T32AG071444 (D.J.A.), F31AG074673 (M.D.T.), T32AG071444 (H.R.S.W.), F30AG079580 (H.R.S.W.), F31AG074628 (T.J.M.), T32DK064466 (D.C.S.), Paul and Carole Stark Medical Neuroscience Fellowship (D.J.A.), Sarah Roush Memorial Fellowship (H.K.), and Eli Lilly-Stark Neuroscience Fellowship (S.P. and L.C.D.). The JK laboratory was also supported by the Strategic Research Initiative (Indiana University) and Indiana University Precision Health Initiative. We thank Dr. Leonard Petrucelli (Mayo Clinic) for providing AAV hTauP301L (RF1 AG062077). This work was also supported, in part, by the Indiana University Pervasive Technology Institute (supported in part by the Lily Endowment Inc.), Shared University Research grants from IBM inc., to Indiana University, the Histology Core of the Indiana Center for Musculoskeletal Health at IU School of Medicine, and the Bone and Body Composition Core of the Indiana Clinical Translational Sciences Institute (CTSI). Sequencing analysis was carried out in the Center for Medical Genomics at Indiana University School of Medicine, which is partially supported by the Indiana University Grand Challenges Precision Health Initiative. We would like to especially acknowledge Windy Woodford, Kari McClimon, and Zachary Bault from the Indiana University Laboratory Animal Resource Center for their assistance with the husbandry of wild-derived mice. The authors declare no competing financial interests. *Author contributions*: D.J.A. and J.K. designed the study and interpreted the results. D.J.A., M.D.T., B.M., A.D.S., S.J., H.K., B.K. T.J.M, D.C.S., S.J.B., and B.T.L. performed the experiments. Y.Y. and C.L.R. performed the tau seeding assay. D.J.A., L.C.D, S.P., and H.R.S.W analyzed the transcriptomic data. D.J.A. and J.K. drafted the manuscript. All authors revised and approved the final manuscript.

## Abbreviations

AAV: Adeno-Associated Virus
AD: Alzheimer’s Disease
AMP-AD: Accelerating Medicines Partnership® Program for Alzheimer’s Disease
B6: C57BL/6J
CAST: CAST/EiJ
DAM: Disease-Associated Microglia
eGFP: enhanced Green Fluorescent Protein
FRET: Fluorescence Resonance Energy Transfer
MAPT: Microtubule Associated Protein Tau
PWK: PWK/PhJ
RD: Repeat Domain
SNP: Single Nucleotide Polymorphism
WSB: WSB/EiJ

## Supplemental Figures

Supplemental Figure 1 **Transcriptomic Analyses for Discovery of Core Tau Response (Signature A)**. Volcano plot demonstrates the log2 fold change (x-axis) and statistical significance (Benjamini-Hochberg adjust p-value, y-axis). (A) B6, (B) CAST, (C) PWK, and (D) WSB mice were analyzed separately to compare genes up-regulated (red, FC > 1.5, p_adj < 0.05) and down-regulated (blue, FC < 1.5, p_adj < 0.05) in Tau-injected mice compared to GFP-injected controls (n = 32/injection group). (E) KEGG Enrichment of Signature A defined in Figure 2C. (F) Heatmap of all Signature A genes in KEGG map05022.

Supplemental Figure 2. **Tau Pathology in hTauP301 expressing wild-derived mice**. (A) Design of a pilot study to determine the sample size. One litter of B6 and WSB mice were injected with AAV-hTauP301L and aged 6 weeks. TBS-soluble protein lysate from the cortex of each pup was transfected into Tau biosensor cells. 24 hours after transfection, cells were trypsanized and FRET+ signal was measured via FACS as a proxy for tau seeding activity. (B) Our pilot study suggests genetic background affects tau seeding activity (nB6 = 6, nWSB = 5; Welch’s t-test *p* = 0.0172). (C) Power analysis based on the seeding activity pilot study suggests a sample size of at least 8 to properly power the main study. (D) Design of the main study to investigate the role of wild-derived mouse genetic background on tauopathy. (E) Representative images show widespread expression of human Tau (HT7+ stain) in injected mice compared to sham-injected control. Images taken at 4X magnification. (F) Representative images suggest the presence of tau aggregates (AT180+ stain) in the cortex of B6 and wild-derived mice injected with AAV-hTauP301L. Images at 20X magnification were taken of the cortex directly superior to the hippocampus at approximately Bregman −1.46mm. (G) Representative Western blot show human Tau (HT7+ blot) in the cortex and hippocampus of AAV-hTauP301L injected mice compared to AAV-eGFP injected controls. (H) Tau protein quantification of total Tau (tTau) and pTau231 (normalized to tTau) show no effect of genetic background. (I) Tau seeding activity of protein extracted from the hippocampus shows no effect of genetic background. (J) Technical replication of tau seeding activity (n = 2 technical replicates per mouse) shows a high correlation in the cortex (R^2^ = 0.7988) and hippocampus (R^2^ = 0.8701).

Supplemental Figure 3. **Summary of Weighted Gene Co-expression Network Analysis (WGCNA)**. Gene module detection was performed using Weighted Gene Co-expression Network Analysis (WGCNA) from hTauP301L-injected and eGFP-injected mice. (A) Sample dendrogram and trait heatmap reveal outlier detection by calculating unbiased sample similarity. Trait heatmap shows samples are segregated out mainly by Injection type and seeding activity score (FRET). (B) Scale independence and mean connectivity calculated by the WGCNA package. Although no thresholds reached the recommended 0.9 threshold for scale free topology, a threshold of 6 was selected based on recommendations of the package’s authors for unsigned network detection in an experiment with at least 40 samples. (C) Module discovery was performed by clustering genes based on topological overlap matrix (TOM) dissimilarity (y-axis: height). Similar clusters were merged using a dissimilarity threshold of 0.25 (merged dynamic). 60 remaining clusters were assigned arbritrary names using R’s color palette. (D) To prioritize modules of interest, quantified traits were correlated to each module’s eigengene expression. Pearson’s R and Benjamini-Hochberg adjusted p-value were reported in each cell of the heatmap (colored by Pearson’s R). Injection defined as binary trait (1: tau, 0: GFP). Genetic background defined as a binary trait (1: wild-derived, 0:B6). Sex defined as a binary trait (1: female, 0: male). FRET defined as a measurement of %Cells with FRET+ signal from Figure 4C).

Supplemental File 1. **Genotyped Variants of Wild-Derived Mice in AMP-AD Nominated Genes**. (A) List of AMP-AD Nominated Targets. Downloaded from Agora 2021/03/05. (B) Summary of genotyped markers in AMP-AD nominated targets. (C) Genotyped variants of wild-derived mice in AMP-AD nominated targets.

Supplemental File 2 **Summary of Genetic Background-Specific Transcriptomic Analyses**. (A) Differential expression analysis of B6.Tau relative to B6.eGFP injected mice (whole B6-transcriptome analysis = 17,963 genes). (B) Differential expression analysis of CAST.Tau relative to CAST.eGFP injected mice (whole CAST-transcriptome analysis = 22,426 genes). (C) Differential expression analysis of PWK.Tau relative to PWK.eGFP injected mice (whole PWK-transcriptome analysis = 22,100 genes). (D) Differential expression analysis of WSB.Tau relative to WSB.eGFP injected mice (whole WSB-transcriptome analysis = 22,399 genes). (E) Upset plot intersection list. Only genes that were measured in each genetic background were kept in this analysis (n = 15,311). Instructions on how to search for specific intersections are found in file. (F) Differential expression analysis with CAST as an interaction term (Tx_Tau_CAST) reveal a unique tau-response not found in B6 mice. (G) Differential expression analysis with PWK as an interaction term (Tx_Tau_PWK) reveal a unique tau-response not found in B6 mice. (H) Differential expression analysis with WSB as an interaction term (Tx_Tau_WSB) reveal a unique tau-response not found in B6 mice. (I) Summary of WGCNA by gene name. This includes the module to which each gene was assigned, the gene significance and p-value for an association to FRET value, gene module membership value, and MMP value. Enrichment analysis output from gprofiler2 for (J) Signature A: core tau-response (K) Signature B: unique tau-response and (L) Signature C: tau-seeding associated response. (M) Sequencing and (N-Q) mapping statistics are summarized for each sample.

## REFERENCES

Arriagada, P. V., Growdon, J. H., Hedley-Whyte, E. T. & Hyman, B. T. 1992. Neurofibrillary tangles but not senile plaques parallel duration and severity of Alzheimer’s disease. Neurology, 42,631–9.

Bailey, R. M., Howard, J., Knight, J., Sahara, N., Dickson, D. W. & Lewis, J. 2014. Effects of the C57BL/6 strain background on tauopathy progression in the rTg4510 mouse model. Mol Neurodegener, 9,8.

Bengoa-Vergniory, N., Velentza-Almpani, E., Silva, A. M., Scott, C., Vargas-Caballero, M., Sastre, M., Wade-Martins, R. & Alegre-Abarrategui, J. 2021. Tau-proximity ligation assay reveals extensive previously undetected pathology prior to neurofibrillary tangles in preclinical Alzheimer’s disease. Acta Neuropathol Commun, 9,18.

Blake, J. A., Baldarelli, R., Kadin, J. A., Richardson, J. E., Smith, C. L., Bult, C. J. & Mouse Genome Database, G. 2021. Mouse Genome Database (MGD): Knowledgebase for mouse-human comparative biology. Nucleic Acids Res, 49,D981–D987.

Carlomagno, Y., Chung, D. C., Yue, M., Kurti, A., Avendano, N. M., Castanedes-Casey, M., Hinkle, K. M., Jansen-West, K., Daughrity, L. M., Tong, J., Phillips, V., Rademakers, R., Deture, M., Fryer, J. D., Dickson, D. W., Petrucelli, L. & Cook, C. 2019. Enhanced phosphorylation of T153 in soluble tau is a defining biochemical feature of the A152T tau risk variant. Acta Neuropathol Commun, 7,10.

Castanho, I., Murray, T. K., Hannon, E., Jeffries, A., Walker, E., Laing, E., Baulf, H., Harvey, J., Bradshaw, L., Randall, A., Moore, K., O’Neill, P., Lunnon, K., Collier, D. A., Ahmed, Z., O’Neill, M. J. & Mill, J. 2020. Transcriptional Signatures of Tau and Amyloid Neuropathology. Cell Rep, 30,2040–2054 e5.

Churchill, G. A., Airey, D. C., Allayee, H., Angel, J. M., Attie, A. D., Beatty, J., Beavis, W. D., Belknap, J. K., Bennett, B., Berrettini, W., Bleich, A., Bogue, M., Broman, K. W., Buck, K. J., Buckler, E., Burmeister, M., Chesler, E. J., Cheverud, J. M., Clapcote, S., Cook, M. N., Cox, R. D., Crabbe, J. C., Crusio, W. E., Darvasi, A., Deschepper, C. F., Doerge, R. W., Farber, C. R., Forejt, J., Gaile, D., Garlow, S. J., Geiger, H., Gershenfeld, H., Gordon, T., Gu, J., Gu, W., De Haan, G., Hayes, N. L., Heller, C., Himmelbauer, H., Hitzemann, R., Hunter, K., Hsu, H. C., Iraqi, F. A., Ivandic, B., Jacob, H. J., Jansen, R. C., Jepsen, K. J., Johnson, D. K., Johnson, T. E., Kempermann, G., Kendziorski, C., Kotb, M., Kooy, R. F., Llamas, B., Lammert, F., Lassalle, J. M., Lowenstein, P. R., Lu, L., Lusis, A., Manly, K. F., Marcucio, R., Matthews, D., Medrano, J. F., Miller, D. R., Mittleman, G., Mock, B. A., Mogil, J. S., Montagutelli, X., Morahan, G., Morris, D. G., Mott, R., Nadeau, J. H., Nagase, H., Nowakowski, R. S., O’Hara, B. F., Osadchuk, A. V., Page, G. P., Paigen, B., Paigen, K., Palmer, A. A., Pan, H. J., Peltonen-Palotie, L., Peirce, J., Pomp, D., Pravenec, M., Prows, D. R., Qi, Z., Reeves, R. H., Roder, J., Rosen, G. D., Schadt, E. E., Schalkwyk, L. C., Seltzer, Z., Shimomura, K., Shou, S., Sillanpaa, M. J., Siracusa, L. D., Snoeck, H. W., Spearow, J. L., Svenson, K., et al. 2004. The Collaborative Cross, a community resource for the genetic analysis of complex traits. Nat Genet, 36,1133–7.

Churchill, G. A., Gatti, D. M., Munger, S. C. & Svenson, K. L. 2012. The Diversity Outbred mouse population. Mamm Genome, 23,713–8.

Clavaguera, F., Bolmont, T., Crowther, R. A., Abramowski, D., Frank, S., Probst, A., Fraser, G., Stalder, A. K., Beibel, M., Staufenbiel, M., Jucker, M., Goedert, M. & Tolnay, M. 2009. Transmission and spreading of tauopathy in transgenic mouse brain. Nat Cell Biol, 11,909–13.

Congdon, E. E. & Sigurdsson, E. M. 2018. Tau-targeting therapies for Alzheimer disease. Nat Rev Neurol, 14,399–415.

Cook, C., Dunmore, J. H., Murray, M. E., Scheffel, K., Shukoor, N., Tong, J., Castanedes-Casey, M., Phillips, V., Rousseau, L., Penuliar, M. S., Kurti, A., Dickson, D. W., Petrucelli, L. & Fryer, J. D. 2014. Severe amygdala dysfunction in a MAPT transgenic mouse model of frontotemporal dementia. Neurobiol Aging, 35,1769–77.

Cook, C., Kang, S. S., Carlomagno, Y., Lin, W. L., Yue, M., Kurti, A., Shinohara, M., Jansen-West, K., Perkerson, E., Castanedes-Casey, M., Rousseau, L., Phillips, V., Bu, G., Dickson, D. W., Petrucelli, L. & Fryer, J. D. 2015. Tau deposition drives neuropathological, inflammatory and behavioral abnormalities independently of neuronal loss in a novel mouse model. Hum Mol Genet, 24,6198–212.

De Calignon, A., Polydoro, M., Suarez-Calvet, M., William, C., Adamowicz, D. H., Kopeikina, K. J., Pitstick, R., Sahara, N., Ashe, K. H., Carlson, G. A., Spires-Jones, T. L. & Hyman, B. T. 2012. Propagation of tau pathology in a model of early Alzheimer’s disease. Neuron, 73,685–97.

Devos, S. L., Corjuc, B. T., Oakley, D. H., Nobuhara, C. K., Bannon, R. N., Chase, A., Commins, C., Gonzalez, J. A., Dooley, P. M., Frosch, M. P. & Hyman, B. T. 2018. Synaptic Tau Seeding Precedes Tau Pathology in Human Alzheimer’s Disease Brain. Front Neurosci, 12,267.

Dujardin, S., Fernandes, A., Bannon, R., Commins, C., De Los Santos, M., Kamath, T. V., Hayashi, M. & Hyman, B. T. 2022. Tau propagation is dependent on the genetic background of mouse strains. Brain Commun, 4,fcac048.

Efthymiou, A. G. & Goate, A. M. 2017. Late onset Alzheimer’s disease genetics implicates microglial pathways in disease risk. Mol Neurodegener, 12,43.

Eskandari-Sedighi, G., Daude, N., Gapeshina, H., Sanders, D. W., Kamali-Jamil, R., Yang, J., Shi, B., Wille, H., Ghetti, B., Diamond, M. I., Janus, C. & Westaway, D. 2017. The CNS in inbred transgenic models of 4-repeat Tauopathy develops consistent tau seeding capacity yet focal and diverse patterns of protein deposition. Mol Neurodegener, 12,72.

Ezerskiy, L. A., Schoch, K. M., Sato, C., Beltcheva, M., Horie, K., Rigo, F., Martynowicz, R., Karch, C. M., Bateman, R. J. & Miller, T. M. 2022. Astrocytic 4R tau expression drives astrocyte reactivity and dysfunction. JCI Insight, 7.

Frost, B., Jacks, R. L. & Diamond, M. I. 2009. Propagation of tau misfolding from the outside to the inside of a cell. J Biol Chem, 284,12845–52.

Gan, K. J., Akram, A., Blasius, T. L., Ramser, E. M., Budaitis, B. G., Gabrych, D. R., Verhey, K. J. & Silverman, M. A. 2020. GSK3beta Impairs KIF1A Transport in a Cellular Model of Alzheimer’s Disease but Does Not Regulate Motor Motility at S402. eNeuro, 7.

He, T., Itano, M. S., Earley, L. F., Hall, N. E., Riddick, N., Samulski, R. J. & Li, C. 2019. The Influence of Murine Genetic Background in Adeno-Associated Virus Transduction of the Mouse Brain. Hum Gene Ther Clin Dev, 30,169–181.

Holmes, B. B., Furman, J. L., Mahan, T. E., Yamasaki, T. R., Mirbaha, H., Eades, W. C., Belaygorod, L., Cairns, N. J., Holtzman, D. M. & Diamond, M. I. 2014. Proteopathic tau seeding predicts tauopathy in vivo. Proc Natl Acad Sci U S A, 111,E4376–85.

Hutton, M., Lendon, C. L., Rizzu, P., Baker, M., Froelich, S., Houlden, H., Pickering-Brown, S., Chakraverty, S., Isaacs, A., Grover, A., Hackett, J., Adamson, J., Lincoln, S., Dickson, D., Davies, P., Petersen, R. C., Stevens, M., De Graaff, E., Wauters, E., Van Baren, J., Hillebrand, M., Joosse, M., Kwon, J. M., Nowotny, P., Che, L. K., Norton, J., Morris, J. C., Reed, L. A., Trojanowski, J., Basun, H., Lannfelt, L., Neystat, M., Fahn, S., Dark, F., Tannenberg, T., Dodd, P. R., Hayward, N., Kwok, J. B., Schofield, P. R., Andreadis, A., Snowden, J., Craufurd, D., Neary, D., Owen, F., Oostra, B. A., Hardy, J., Goate, A., Van Swieten, J., Mann, D., Lynch, T. & Heutink, P. 1998. Association of missense and 5’-splice-site mutations in tau with the inherited dementia FTDP-17. Nature, 393,702–5.

Jin, N., Gu, J., Wu, R., Chu, D., Tung, Y. C., Wegiel, J., Wisniewski, T., Gong, C. X., Iqbal, K. & Liu, F. 2022. Tau seeding activity in various regions of down syndrome brain assessed by two novel assays. Acta Neuropathol Commun, 10,132.

Kang, S. S., Ebbert, M. T. W., Baker, K. E., Cook, C., Wang, X., Sens, J. P., Kocher, J. P., Petrucelli, L. & Fryer, J. D. 2018. Microglial translational profiling reveals a convergent APOE pathway from aging, amyloid, and tau. J Exp Med, 215,2235–2245.

Karahan, H., Smith, D. C., Kim, B., Dabin, L. C., Al-Amin, M. M., Wijeratne, H. R. S., Pennington, T., Viana Di Prisco, G., Mccord, B., Lin, P. B., Li, Y., Peng, J., Oblak, A. L., Chu, S., Atwood, B. K. & Kim, J. 2021. Deletion of Abi3 gene locus exacerbates neuropathological features of Alzheimer’s disease in a mouse model of Abeta amyloidosis. Sci Adv, 7,eabe3954.

Karch, C. M. & Goate, A. M. 2015. Alzheimer’s disease risk genes and mechanisms of disease pathogenesis. Biol Psychiatry, 77,43–51.

Kaufman, S. K., Del Tredici, K., Thomas, T. L., Braak, H. & Diamond, M. I. 2018. Tau seeding activity begins in the transentorhinal/entorhinal regions and anticipates phospho-tau pathology in Alzheimer’s disease and PART. Acta Neuropathol, 136,57–67.

Keren-Shaul, H., Spinrad, A., Weiner, A., Matcovitch-Natan, O., Dvir-Szternfeld, R., Ulland, T. K., David, E., Baruch, K., Lara-Astaiso, D., Toth, B., Itzkovitz, S., Colonna, M., Schwartz, M. & Amit, I. 2017. A Unique Microglia Type Associated with Restricting Development of Alzheimer’s Disease. Cell, 169,1276–1290 e17.

Kim, J., Miller, V. M., Levites, Y., West, K. J., Zwizinski, C. W., Moore, B. D., Troendle, F. J., Bann, M., Verbeeck, C., Price, R. W., Smithson, L., Sonoda, L., Wagg, K., Rangachari, V., Zou, F., Younkin, S. G., Graff-Radford, N., Dickson, D., Rosenberry, T. & Golde, T. E. 2008. BRI2 (ITM2b) inhibits Abeta deposition in vivo. J Neurosci, 28,6030–6.

Kim, J. Y., Grunke, S. D., Levites, Y., Golde, T. E. & Jankowsky, J. L. 2014. Intracerebroventricular viral injection of the neonatal mouse brain for persistent and widespread neuronal transduction. J Vis Exp,51863.

Kollmus, H., Fuchs, H., Lengger, C., Haselimashhadi, H., Bogue, M. A., Ostereicher, M. A., Horsch, M., Adler, T., Aguilar-Pimentel, J. A., Amarie, O. V., Becker, L., Beckers, J., Calzada-Wack, J., Garrett, L., Hans, W., Holter, S. M., Klein-Rodewald, T., Maier, H., Mayer-Kuckuk, P., Miller, G., Moreth, K., Neff, F., Rathkolb, B., Racz, I., Rozman, J., Spielmann, N., Treise, I., Busch, D., Graw, J., Klopstock, T., Wolf, E., Wurst, W., Yildirim, A. O., Mason, J., Torres, A., MOUSE PHENOME DATABASE, T., Balling, R., Mehaan, T., Gailus-Durner, V., Schughart, K. & Hrabe De Angelis, M. 2020. A comprehensive and comparative phenotypic analysis of the collaborative founder strains identifies new and known phenotypes. Mamm Genome, 31,30–48.

Kraus, A., Saijo, E., Metrick, M. A., 2nd, Newell, K., Sigurdson, C. J., Zanusso, G., Ghetti, B. & Caughey, B. 2019. Seeding selectivity and ultrasensitive detection of tau aggregate conformers of Alzheimer disease. Acta Neuropathol, 137,585–598.

Langfelder, P. & Horvath, S. 2008. WGCNA: an R package for weighted correlation network analysis. BMC Bioinformatics, 9,559.

Lasagna-Reeves, C. A., De Haro, M., Hao, S., Park, J., Rousseaux, M. W., Al-Ramahi, I., Jafar-Nejad, P., Vilanova-Velez, L., See, L., De Maio, A., Nitschke, L., Wu, Z., Troncoso, J. C., Westbrook, T. F., Tang, J., Botas, J. & Zoghbi, H. Y. 2016. Reduction of Nuak1 Decreases Tau and Reverses Phenotypes in a Tauopathy Mouse Model. Neuron, 92,407–418.

Lee, V. M., Brunden, K. R., Hutton, M. & Trojanowski, J. Q. 2011. Developing therapeutic approaches to tau, selected kinases, and related neuronal protein targets. Cold Spring Harb Perspect Med, 1,a006437.

Liao, Y., Smyth, G. K. & Shi, W. 2014. featureCounts: an efficient general purpose program for assigning sequence reads to genomic features. Bioinformatics, 30,923–30.

Lilue, J., Doran, A. G., Fiddes, I. T., Abrudan, M., Armstrong, J., Bennett, R., Chow, W., Collins, J., Collins, S., Czechanski, A., Danecek, P., Diekhans, M., Dolle, D. D., Dunn, M., Durbin, R., Earl, D., Ferguson-Smith, A., Flicek, P., Flint, J., Frankish, A., Fu, B., Gerstein, M., Gilbert, J., Goodstadt, L., Harrow, J., Howe, K., Ibarra-Soria, X., Kolmogorov, M., Lelliott, C. J., Logan, D. W., Loveland, J., Mathews, C. E., Mott, R., Muir, P., Nachtweide, S., Navarro, F. C. P., Odom, D. T., Park, N., Pelan, S., Pham, S. K., Quail, M., Reinholdt, L., Romoth, L., Shirley, L., Sisu, C., Sjoberg-Herrera, M., Stanke, M., Steward, C., Thomas, M., Threadgold, G., Thybert, D., Torrance, J., Wong, K., Wood, J., Yalcin, B., Yang, F., Adams, D. J., Paten, B. & Keane, T. M. 2018. Sixteen diverse laboratory mouse reference genomes define strain-specific haplotypes and novel functional loci. Nat Genet, 50,1574–1583.

Long, J. M. & Holtzman, D. M. 2019. Alzheimer Disease: An Update on Pathobiology and Treatment Strategies. Cell, 179,312–339.

Love, M. I., Huber, W. & Anders, S. 2014. Moderated estimation of fold change and dispersion for RNA-seq data with DESeq2. Genome Biol, 15,550.

Makino, S., Kunimoto, K., Muraoka, Y., Mizushima, Y., Katagiri, K. & Tochino, Y. 1980. Breeding of a non-obese, diabetic strain of mice. Jikken Dobutsu, 29,1–13.

Martinez, P., Patel, H., You, Y., Jury, N., Perkins, A., Lee-Gosselin, A., Taylor, X., You, Y., Viana Di Prisco, G., Huang, X., Dutta, S., Wijeratne, A. B., Redding-Ochoa, J., Shahid, S. S., Codocedo, J. F., Min, S., Landreth, G. E., Mosley, A. L., Wu, Y. C., Mckinzie, D. L., Rochet, J. C., Zhang, J., Atwood, B. K., Troncoso, J. & Lasagna-Reeves, C. A. 2022. Bassoon contributes to tau-seed propagation and neurotoxicity. Nat Neurosci, 25,1597–1607.

Mekada, K., Abe, K., Murakami, A., Nakamura, S., Nakata, H., Moriwaki, K., Obata, Y. & Yoshiki, A.2009. Genetic differences among C57BL/6 substrains. Exp Anim, 58,141–9.

Mhatre, S. D., Tsai, C. A., Rubin, A. J., James, M. L. & Andreasson, K. I. 2015. Microglial malfunction: the third rail in the development of Alzheimer’s disease. Trends Neurosci, 38,621–636.

Mirbaha, H., Chen, D., Mullapudi, V., Terpack, S. J., White, C. L., 3RD, Joachimiak, L. A. & Diamond, M. I. 2022. Seed-competent tau monomer initiates pathology in a tauopathy mouse model. J Biol Chem, 298,102163.

Monzon-Sandoval, J., Burlacu, E., Agarwal, D., Handel, A. E., Wei, L., Davis, J., Cowley, S. A., Cader, M. Z. & Webber, C. 2022. Lipopolysaccharide distinctively alters human microglia transcriptomes to resemble microglia from Alzheimer’s disease mouse models. Dis Model Mech, 15.

Morgan, A. P. 2015. argyle: An R Package for Analysis of Illumina Genotyping Arrays. G3 (Bethesda), 6,281–6.

Morgan, A. P., Fu, C. P., Kao, C. Y., Welsh, C. E., Didion, J. P., Yadgary, L., Hyacinth, L., Ferris, M. T., Bell, T. A., Miller, D. R., Giusti-Rodriguez, P., Nonneman, R. J., Cook, K. D., Whitmire, J. K., Gralinski, L. E., Keller, M., Attie, A. D., Churchill, G. A., Petkov, P., Sullivan, P. F., Brennan, J. R., Mcmillan, L. & Pardo-Manuel De Villena, F. 2015. The Mouse Universal Genotyping Array: From Substrains to Subspecies. G3 (Bethesda), 6,263–79.

Mouse Genome Sequencing, C., Waterston, R. H., Lindblad-Toh, K., Birney, E., Rogers, J., Abril, J. F., Agarwal, P., Agarwala, R., Ainscough, R., Alexandersson, M., An, P., Antonarakis, S. E., Attwood, J., Baertsch, R., Bailey, J., Barlow, K., Beck, S., Berry, E., Birren, B., Bloom, T., Bork, P., Botcherby, M., Bray, N., Brent, M. R., Brown, D. G., Brown, S. D., Bult, C., Burton, J., Butler, J., Campbell, R. D., Carninci, P., Cawley, S., Chiaromonte, F., Chinwalla, A. T., Church, D. M., Clamp, M., Clee, C., Collins, F. S., Cook, L. L., Copley, R. R., Coulson, A., Couronne, O., Cuff, J., Curwen, V., Cutts, T., Daly, M., David, R., Davies, J., Delehaunty, K. D., Deri, J., Dermitzakis, E. T., Dewey, C., Dickens, N. J., Diekhans, M., Dodge, S., Dubchak, I., Dunn, D. M., Eddy, S. R., Elnitski, L., Emes, R. D., Eswara, P., Eyras, E., Felsenfeld, A., Fewell, G. A., Flicek, P., Foley, K., Frankel, W. N., Fulton, L. A., Fulton, R. S., Furey, T. S., Gage, D., Gibbs, R. A., Glusman, G., Gnerre, S., Goldman, N., Goodstadt, L., Grafham, D., Graves, T. A., Green, E. D., Gregory, S., Guigo, R., Guyer, M., Hardison, R. C., Haussler, D., Hayashizaki, Y., Hillier, L. W., Hinrichs, A., Hlavina, W., Holzer, T., Hsu, F., Hua, A., Hubbard, T., Hunt, A., Jackson, I., Jaffe, D. B., Johnson, L. S., Jones, M., Jones, T. A., Joy, A., Kamal, M., et al. 2002. Initial sequencing and comparative analysis of the mouse genome. Nature, 420,520–62.

Onos, K. D., Quinney, S. K., Jones, D. R., Masters, A. R., Pandey, R., Keezer, K. J., Biesdorf, C., Metzger, I. F., Meyers, J. A., Peters, J., Persohn, S. C., Mccarthy, B. P., Bedwell, A. A., Figueiredo, L. L., Cope, Z. A., Sasner, M., Howell, G. R., Williams, H. M., Oblak, A. L., Lamb, B. T., Carter, G. W., Rizzo, S. J. S. & Territo, P. R. 2022. Pharmacokinetic, pharmacodynamic, and transcriptomic analysis of chronic levetiracetam treatment in 5XFAD mice: A MODEL-AD preclinical testing core study. Alzheimers Dement (N Y), 8,e12329

Onos, K. D., Uyar, A., Keezer, K. J., Jackson, H. M., Preuss, C., Acklin, C. J., O’Rourke, R., Buchanan, R., Cossette, T. L., Sukoff Rizzo, S. J., Soto, I., Carter, G. W. & Howell, G. R. 2019. Enhancing face validity of mouse models of Alzheimer’s disease with natural genetic variation. PLoS Genet, 15,e1008155.

Paolicelli, R. C., Sierra, A., Stevens, B., Tremblay, M. E., Aguzzi, A., Ajami, B., Amit, I., Audinat, E., Bechmann, I., Bennett, M., Bennett, F., Bessis, A., Biber, K., Bilbo, S., Blurton-Jones, M., Boddeke, E., Brites, D., Brone, B., Brown, G. C., Butovsky, O., Carson, M. J., Castellano, B., Colonna, M., Cowley, S. A., Cunningham, C., Davalos, D., De Jager, P. L., De Strooper, B., Denes, A., Eggen, B. J. L., Eyo, U., Galea, E., Garel, S., Ginhoux, F., Glass, C. K., Gokce, O., Gomez-Nicola, D., Gonzalez, B., Gordon, S., Graeber, M. B., Greenhalgh, A. D., Gressens, P., Greter, M., Gutmann, D. H., Haass, C., Heneka, M. T., Heppner, F. L., Hong, S., Hume, D. A., Jung, S., Kettenmann, H., Kipnis, J., Koyama, R., Lemke, G., Lynch, M., Majewska, A., Malcangio, M., Malm, T., Mancuso, R., Masuda, T., Matteoli, M., Mccoll, B. W., Miron, V. E., Molofsky, A. V., Monje, M., Mracsko, E., Nadjar, A., Neher, J. J., Neniskyte, U., Neumann, H., Noda, M., Peng, B., Peri, F., Perry, V. H., Popovich, P. G., Pridans, C., Priller, J., Prinz, M., Ragozzino, D., Ransohoff, R. M., Salter, M. W., Schaefer, A., Schafer, D. P., Schwartz, M., Simons, M., Smith, C. J., Streit, W. J., Tay, T. L., Tsai, L. H., Verkhratsky, A., Von Bernhardi, R., Wake, H., Wittamer, V., Wolf, S. A., Wu, L. J. & Wyss-Coray, T. 2022. Microglia states and nomenclature: A field at its crossroads. Neuron, 110,3458–3483.

Passini, M. A., Watson, D. J., Vite, C. H., Landsburg, D. J., Feigenbaum, A. L. & Wolfe, J. H. 2003. Intraventricular brain injection of adeno-associated virus type 1 (AAV1) in neonatal mice results in complementary patterns of neuronal transduction to AAV2 and total long-term correction of storage lesions in the brains of beta-glucuronidase-deficient mice. J Virol, 77,7034–40.

Peirce, J. L., Lu, L., Gu, J., Silver, L. M. & Williams, R. W. 2004. A new set of BXD recombinant inbred lines from advanced intercross populations in mice. BMC Genet, 5,7.

Peng, C., Trojanowski, J. Q. & Lee, V. M. 2020. Protein transmission in neurodegenerative disease. Nat Rev Neurol, 16,199–212.

Poorkaj, P., Bird, T. D., Wijsman, E., Nemens, E., Garruto, R. M., Anderson, L., Andreadis, A., Wiederholt, W. C., Raskind, M. & Schellenberg, G. D. 1998. Tau is a candidate gene for chromosome 17 frontotemporal dementia. Ann Neurol, 43,815–25.

Rauch, J. N., Luna, G., Guzman, E., Audouard, M., Challis, C., Sibih, Y. E., Leshuk, C., Hernandez, I., Wegmann, S., Hyman, B. T., Gradinaru, V., Kampmann, M. & Kosik, K. S. 2020. LRP1 is a master regulator of tau uptake and spread. Nature, 580,381–385.

Santacruz, K., Lewis, J., Spires, T., Paulson, J., Kotilinek, L., Ingelsson, M., Guimaraes, A., Deture, M., Ramsden, M., Mcgowan, E., Forster, C., Yue, M., Orne, J., Janus, C., Mariash, A., Kuskowski, M., Hyman, B., Hutton, M. & Ashe, K. H. 2005. Tau suppression in a neurodegenerative mouse model improves memory function. Science, 309,476–81.

Sarsani, V. K., Raghupathy, N., Fiddes, I. T., Armstrong, J., Thibaud-Nissen, F., Zinder, O., Bolisetty, M., Howe, K., Hinerfeld, D., Ruan, X., Rowe, L., Barter, M., Ananda, G., Paten, B., Weinstock, G. M., Churchill, G. A., Wiles, M. V., Schneider, V. A., Srivastava, A. & Reinholdt, L. G. 2019. The Genome of C57BL/6J “Eve", the Mother of the Laboratory Mouse Genome Reference Strain. G3 (Bethesda), 9,1795–1805.

Schoch, K. M., Ezerskiy, L. A., Morhaus, M. M., Bannon, R. N., Sauerbeck, A. D., Shabsovich, M., Jafar-Nejad, P., Rigo, F. & Miller, T. M. 2021. Acute Trem2 reduction triggers increased microglial phagocytosis, slowing amyloid deposition in mice. Proc Natl Acad Sci U S A, 118.

Sierksma, A., Lu, A., Mancuso, R., Fattorelli, N., Thrupp, N., Salta, E., Zoco, J., Blum, D., Buee, L., De Strooper, B. & Fiers, M. 2020. Novel Alzheimer risk genes determine the microglia response to amyloid-beta but not to TAU pathology. EMBO Mol Med, 12,e10606.

Simon, M. M., Greenaway, S., White, J. K., Fuchs, H., Gailus-Durner, V., Wells, S., Sorg, T., Wong, K., Bedu, E., Cartwright, E. J., Dacquin, R., Djebali, S., Estabel, J., Graw, J., Ingham, N. J., Jackson, I. J., Lengeling, A., Mandillo, S., Marvel, J., Meziane, H., Preitner, F., Puk, O., Roux, M., Adams, D. J., Atkins, S., Ayadi, A., Becker, L., Blake, A., Brooker, D., Cater, H., Champy, M. F., Combe, R., Danecek, P., Di Fenza, A., Gates, H., Gerdin, A. K., Golini, E., Hancock, J. M., Hans, W., Holter, S. M., Hough, T., Jurdic, P., Keane, T. M., Morgan, H., Muller, W., Neff, F., Nicholson, G., Pasche, B., Roberson, L. A., Rozman, J., Sanderson, M., Santos, L., Selloum, M., Shannon, C., Southwell, A., Tocchini-Valentini, G. P., Vancollie, V. E., Westerberg, H., Wurst, W., Zi, M., Yalcin, B., Ramirez-Solis, R., Steel, K. P., Mallon, A. M., De Angelis, M. H., Herault, Y. & Brown, S. D. 2013. A comparative phenotypic and genomic analysis of C57BL/6J and C57BL/6N mouse strains. Genome Biol, 14,R82.

Stopschinski, B. E., Del Tredici, K., Estill-Terpack, S. J., Ghebremdehin, E., Yu, F. F., Braak, H. & Diamond, M. I. 2021. Anatomic survey of seeding in Alzheimer’s disease brains reveals unexpected patterns. Acta Neuropathol Commun, 9,164.

Wegmann, S., Bennett, R. E., Amaral, A. S. & Hyman, B. T. 2017. Studying tau protein propagation and pathology in the mouse brain using adeno-associated viruses. Methods Cell Biol, 141,307–322.

Woerman, A. L., Patel, S., Kazmi, S. A., Oehler, A., Freyman, Y., Espiritu, L., Cotter, R., Castaneda, J. A., Olson, S. H. & Prusiner, S. B. 2017. Kinetics of Human Mutant Tau Prion Formation in the Brains of 2 Transgenic Mouse Lines. JAMA Neurol, 74,1464–1472.

Yalcin, B., Wong, K., Agam, A., Goodson, M., Keane, T. M., Gan, X., Nellaker, C., Goodstadt, L., Nicod, J., Bhomra, A., Hernandez-Pliego, P., Whitley, H., Cleak, J., Dutton, R., Janowitz, D., Mott, R., Adams, D. J. & Flint, J. 2011. Sequence-based characterization of structural variation in the mouse genome. Nature, 477,326–9.

Yanagisawa, D., Hamezah, H. S., Pahrudin Arrozi, A. & Tooyama, I. 2021. Differential accumulation of tau pathology between reciprocal F1 hybrids of rTg4510 mice. Sci Rep, 11,9623.

Yang, H., Wang, J. R., Didion, J. P., Buus, R. J., Bell, T. A., Welsh, C. E., Bonhomme, F., Yu, A. H., Nachman, M. W., Pialek, J., Tucker, P., Boursot, P., Mcmillan, L., Churchill, G. A. & De Villena, F. P. 2011. Subspecific origin and haplotype diversity in the laboratory mouse. Nat Genet, 43,648–55.

Yang, H. S., Onos, K. D., Choi, K., Keezer, K. J., Skelly, D. A., Carter, G. W. & Howell, G. R. 2021. Natural genetic variation determines microglia heterogeneity in wild-derived mouse models of Alzheimer’s disease. Cell Rep, 34,108739.

Yoshiyama, Y., Higuchi, M., Zhang, B., Huang, S. M., Iwata, N., Saido, T. C., Maeda, J., Suhara, T., Trojanowski, J. Q. & Lee, V. M. 2007. Synapse loss and microglial activation precede tangles in a P301S tauopathy mouse model. Neuron, 53,337–51.

Zhu, A., Ibrahim, J. G. & Love, M. I. 2019. Heavy-tailed prior distributions for sequence count data: removing the noise and preserving large differences. Bioinformatics, 35,2084–2092.

